# Broad receptor tropism and immunogenicity of a clade 3 sarbecovirus

**DOI:** 10.1101/2023.09.12.557371

**Authors:** Jimin Lee, Samantha K. Zepeda, Young-Jun Park, Ashley L. Taylor, Joel Quispe, Cameron Stewart, Elizabeth M. Leaf, Catherine Treichel, Davide Corti, Neil P. King, Tyler N. Starr, David Veesler

## Abstract

Although *Rhinolophus* bats harbor diverse clade 3 sarbecoviruses, the structural determinants of receptor tropism along with the antigenicity of their spike (S) glycoproteins remain uncharacterized. Here, we show that the African Rinolophus bat clade 3 sarbecovirus PRD-0038 S has a broad ACE2 usage and that RBD mutations further expand receptor promiscuity and enable human ACE2 utilization. We determined a cryoEM structure of the PRD-0038 RBD bound to *R. alcyone* ACE2, explaining receptor tropism and highlighting differences with SARS-CoV-1 and SARS-CoV-2. Characterization of PRD-0038 S using cryoEM and monoclonal antibody reactivity revealed its distinct antigenicity relative to SARS-CoV-2 and identified PRD-0038 cross-neutralizing antibodies for pandemic preparedness. PRD-0038 S vaccination elicited greater titers of antibodies cross-reacting with vaccine-mismatched clade 2 and clade 1a sarbecoviruses compared to SARS-CoV-2 S due to broader antigenic targeting, motivating the inclusion of clade 3 antigens in next-generation vaccines for enhanced resilience to viral evolution.

## Introduction

Two sarbecoviruses have crossed the species barrier and spilled over to humans in the past two decades. SARS-CoV-1 emerged in 2002 and spread worldwide through air travel routes, leading to an epidemic with 8,098 cases and 774 deaths^1,2^. SARS-CoV-2 emerged at the end of 2019 and led to the devastating COVID-19 pandemic which claimed millions of lives worldwide^3,4^. Both viruses enter human cells via spike (S)-mediated fusion of the viral and host membranes upon binding to the angiotensin-converting enzyme 2 (ACE2) receptor^3,5–8^.

Reports of additional sarbecovirus spillovers to humans^9,10^ along with detection of numerous sarbecoviruses in bats and other wild animals^3,11–16^ underscore the recurrent zoonotic threat to public health posed by these viruses. The S glycoprotein of some of these sarbecoviruses harbor a receptor-binding domain (RBD) that utilize the human ACE2 receptor to enter host cells, indicating that they could possibly cross the species barrier to infect humans^6,12,17–20^.

Phylogenetic classification of sarbecoviruses based on their RBD sequences led to the definition of at least four clades: clade 1a (e.g. SARS-CoV-1), clade 1b (e.g. SARS-CoV-2), clade 2 (e.g. RmYN02) and clade 3 (e.g. BtKY72) ^21,22^. Clade 3 sarbecoviruses have been identified in bats in Europe and Africa^23–28^, such as BtKY72 and PRD-0038 for which sequences were found in Kenya and Rwanda, respectively. We recently showed that the S glycoprotein of one of them (BtKY72) could utilize two *Rhinolophus affinis* ACE2 alleles to promote entry into cells^29^. Furthermore, two amino acid residue substitutions in the BtKY72 RBD enabled S-mediated entry into human ACE2-expressing cells, broadening the range of sarbecoviruses with spillover potential^29^. The importance of this observation was underscored by the recent discovery of the clade 3 Khosta-2 virus^28^, which independently acquired the ability to bind^29^ and enter cells^30^ using the human ACE2 receptor. Studying the structure and functional properties of clade 3 sarbecovirus spike (S) glycoproteins is therefore crucial to understand spillover risk and assist in pandemic preparedness.

Here, we report that the S glycoprotein of the clade 3 sarbecovirus PRD-0038, which is a member of the largely uncharacterized African bat-borne sarbecoviruses, has a broad ACE2 usage and that PRD-0038 RBD mutations further expand entry receptor tropism to additional *Rhinolophus* bat species and human ACE2. We determined structures of the PRD-0038 RBD bound to *R. alcyone* ACE2 and of the PRD-0038 S trimer, explaining receptor tropism and the distinct antigenicity of clade 3 sarbecoviruses relative to SARS-CoV-2 and SARS-CoV-1. Evaluation of a panel of monoclonal antibodies enabled identification of PRD-0038 cross-neutralizing antibodies that could be deployed for outbreak response. Vaccination of mice with PRD-0038 S elicited greater titers of antibodies cross-reacting with vaccine-mismatched clade 2 and clade 1a sarbecoviruses, relative to SARS-CoV-2 S immunization, indicating that addition of clade 3 antigens in vaccine formulations could enhance the resilience of antibody responses to viral evolution. Our findings highlight a molecular pathway for possible zoonotic spillover of a clade 3 sarbecovirus and the necessity of developing pan-sarbecovirus vaccines and countermeasures.

## RESULTS

### PRD-0038 can utilize a broad spectrum of *Rhinolophus* bat ACE2 orthologs as entry receptors

To investigate the promiscuity of clade 3 sarbecovirus host receptor usage, we first assessed binding of a panel of *Rhinolophus* bat ACE2 orthologs harboring a C-terminal human Fc fusion to the immobilized PRD-0038 RBD using biolayer interferometry (BLI) (**Figures 1A and 1B**). We selected PRD-0038 as a representative member of African bat-borne sarbecoviruses due to its more ancestral phylogenetic positioning relative to the other two sarbecoviruses isolated on the same continent^29^ (BtKY72 and PDF-2370) and the high sequence similarity of their S glycoproteins. Our ACE2 panel comprised eight distinct *R. sinicus* alleles and two distinct *R. affinis* alleles, which were defined based on polymorphisms within the region recognized by sarbecovirus RBDs^29,31^, as well as *R. alcyone* and *R. landeri* orthologues. *R. sinicus* Asian bats are probable reservoir hosts for SARS-CoV-1^14^, *R. affinis* bats have been shown to host closely related viruses to SARS-CoV-2^3^ whereas *R. alcyone* and *R. landeri* bats are found in sub-saharan Africa, overlapping with the regions of sampling of several clade 3 sarbecoviruses (the exact *Rhinolophus* species from which PRD-0038 and BtKY72 have been sampled is unknown)^24,25^ (**Figure 1C**). We observed the strongest binding to the PRD-0038 RBD with *R. alcyone* ACE2, which exhibited the slowest dissociation kinetics in our panel (**Figure 1A**). The PRD-0038 RBD also interacted with both *R. affinis* ACE2s, albeit more tightly with the 9479 allele than the 787 allele, and with *R. landeri* ACE2 (**Figure 1A**). Finally, we detected binding to four out of the eight *R. sinicus* alleles evaluated (**Figure 1B**).

**Figure 1.**
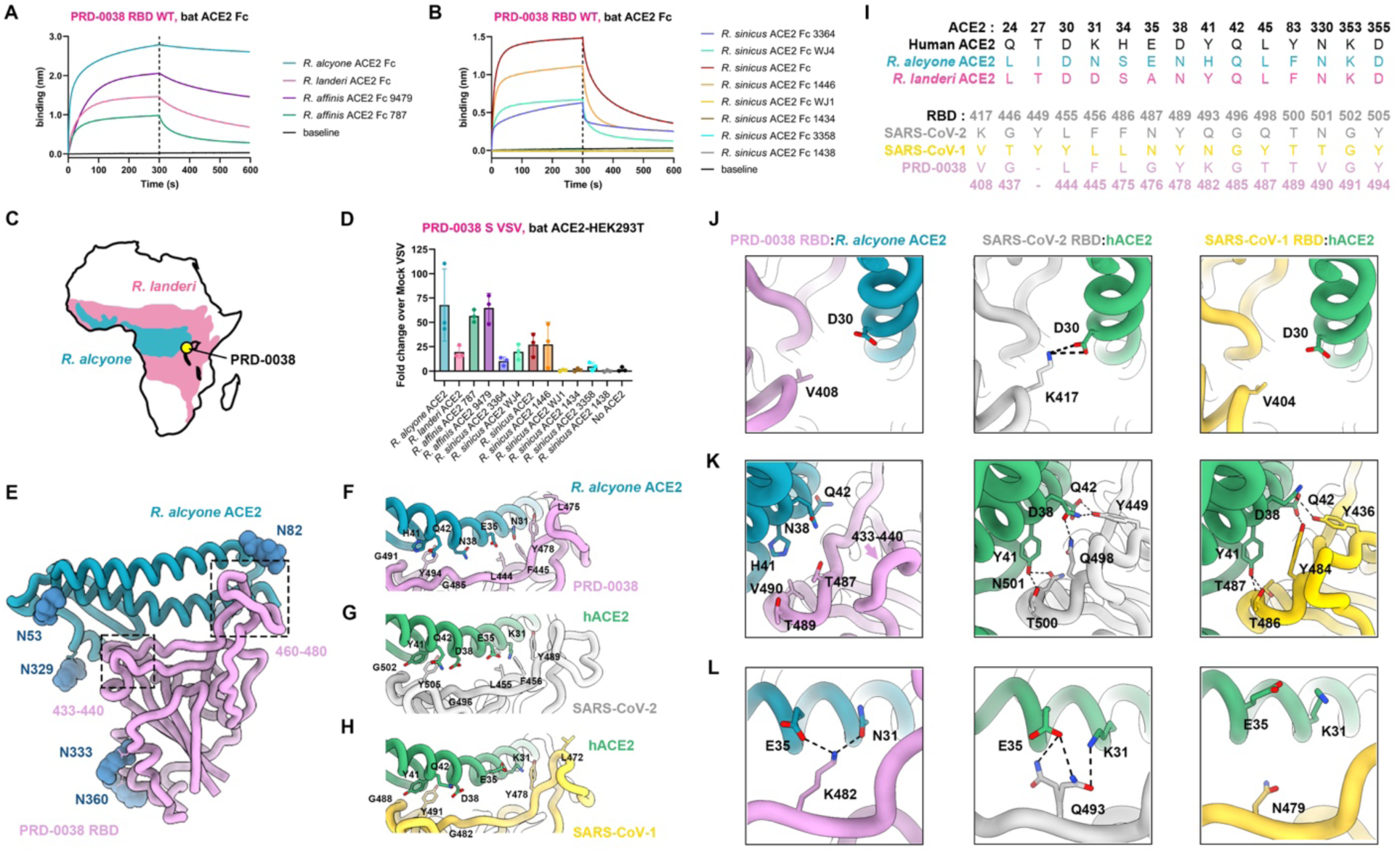
The clade 3 PRD-0038 sarbecovirus has a broad *Rhinolophus* bat ACE2 tropism. (A and B) BLI binding analysis of *R. affinis*, *R. alcyone* and *R. landeri* (see also Table S1) (A) or *R. sinicus* (B) ACE2-Fc alleles at a concentration of 1 µM to the biotinylated PRD-0038 RBD immobilized on streptavidin biosensors. Baselines represent non-specific binding of non-coated streptavidin biosensors to 0.25 µM ACE2 Fc. (C) Known geographic distribution of *R. alcyone* and *R. landeri* bats in sub-saharan Africa (https://www.iucnredlist.org/). The yellow area indicates the site of PRD-0038 sampling (Rwanda). (D) Entry of VSV pseudotyped with wildtype PRD-0038 S in HEK293T cells transiently transfected with the indicated *Rhinolophus* bat ACE2 orthologs. Each point represents the average of technical duplicates from each biological triplicate. Means and standard deviations shown as bars and error bars. (E) CryoEM structure of the PRD-0038 RBD bound to *R. alcyone* ACE2. N-linked glycans are shown as dark blue spheres. The dotted gray boxes highlight regions with major structural deviations from SARS-CoV-2 (see also Figure S2) (F-H) Close-up views of the interface between the PRD-0038 RBD and *R. alcyone* ACE2 (F, pink and blue, respectively), the SARS-CoV-2 RBD and human ACE2 (G, gray and green, respectively, PDB ID 6M0J^33^) and the SARS-CoV-1 RBD and human ACE2 (H, gold and green, respectively, PDB ID 2AJF^34^). Key conserved residues at the interface are rendered as sticks. (I) Sequence alignments of the key ACE2 and RBD residues at the binding interfaces. Numberings used are for human ACE2 and SARS-CoV-2 (see also Table S1). (J-L) Close-up views of selected key contact residues at the interface between the PRD-0038 RBD and *R. alcyone* ACE2, the SARS-CoV-2 RBD and human ACE2 and the SARS-CoV-1 RBD and human ACE2 colored as in panel (F).

To further evaluate receptor tropism, we pseudotyped vesicular stomatitis virus (VSV) particles with PRD-0038 S and assessed entry into HEK293T cells transiently transfected with the corresponding set of full-length, membrane-anchored ACE2s (**Figure 1D**). Concurring with our binding data, *R. alcyone* enabled robust entry of PRD-0038 S VSV (**Figure 1D**), in line with a previous study^32^. Moreover, we also detected efficient entry into cells expressing *R. affinis* 9479 and 787 ACE2 alleles, *R. landeri* ACE2 and the four *R. sinicus* ACE2 alleles for which binding was detected by BLI (**Figure 1D**). Collectively, these data show that PRD-0038 S recognizes and can utilize a broad spectrum of *Rhinolophus* bat ACE2 orthologs as entry receptors, including those from bat species known to be found in geographic areas proximal to the site of PRD-0038 discovery.

### Molecular basis of PRD-0038 RBD engagement of the *R. alcyone* ACE2 receptor

To reveal the structural determinants of ACE2 recognition by clade 3 sarbecoviruses, we determined a cryoEM reconstruction of the PRD-0038 RBD bound to the natively dimeric *R. alcyone* ACE2 (RaACE2) ectodomain using single-particle cryo-electron microscopy (cryoEM) (**Figures 1E, S1, and Table S2**). Symmetry-expansion and local refinement yielded a structure at 3.2 Å resolution of the ACE2 peptidase domain bound to the RBD revealing the molecular interactions mediating complex formation. An average surface of ∼750 Å^2^ is buried at the ACE2/RBD interface as compared to ∼840 Å^2^ for the complexes of human ACE2 (hACE2) bound to the SARS-CoV-2 RBD^33^ or to the SARS-CoV-1 RBD^33,34^. The relative orientation of the binding partners are similar for these three structures, likely due to conservation of several key ACE2-interacting residues including L444_PRD-0038_/L455_SARS-CoV-2_ and F445_PRD-0038_/F456_SARS-CoV-2_, L475_PRD-0038_/L472_SARS-CoV-1_, Y478_PRD-0038_/Y489_SARS-CoV-2_, G485_PRD-0038_/G496_SARS-CoV-2_, G491_PRD-0038_/G502_SARS-CoV-2_ and Y494_PRD-0038_/Y505_SARS-CoV-2_ (**Figures 1F-1H**). However, the salt bridge formed between K417_SARS-CoV-2_ and D30_hACE2_ is absent due to substitution to V408_PRD-0038_ and V404_SARS-CoV-1_ (**Figures 1F, 1G, and 1J**). Furthermore, the electrostatic interactions involving residues D38_hACE2_ and Q42_hACE2_ with Y449_SARS-CoV-2_ and Q498_SARS-CoV-2_ or Y436_SARS-CoV-1_ and Y484_SARS-CoV-1_ are lost are lost due to and remodeling of the PRD-0038 433-440 loop along with substitution of Q498_SARS-CoV-2_ to T487_PRD-0038_ (**Figures 1F-1H, 1K, and S2**). The interface between T500/N501_SARS-CoV-2_ (or Y501 in currently circulating variants) and Y41_hACE2_ is replaced by tenuous contacts between the topologically equivalent residues T489/V490_PRD-0038_ and H41_RaACE2_ (**Figures 1F, 1G, and 1K**). Q493_SARS-CoV-2_ optimally interacts with K31_hACE2_/E35_hACE2_ whereas the topologically equivalent residue K482_PRD-0038_ is better adapted to N31_RaACE2_/E35_RaACE2_ due to swapping of the position of a positively charged amino acid side chain across the interface (**Figures 1F, 1G, and 1L**). These findings concur with (i) the enhanced entry of the closely-related BtKY72 RBD-harboring pseudovirus into cells expressing a human ACE2 K31D mutant relative to wildtype ACE2^32^; (ii) the isolation of a mouse-adapted SARS-CoV-2 isolate harboring the Q493K substitution^35^ that promotes favorable interactions with mouse ACE2, the latter ortholog also possessing residues N31/E35^36,37^; and (iii) the emergence of R493_SARS-CoV-2_ in Omicron BA.1 and BA.2^36,37^, which was subsequently reverted to the more favorable Q493_SARS-CoV-2_ in subsequent variants, likely due to relieving electrostatic repulsion with K31_hACE2_^38^. Overall, most of the PRD-0038 RBD binding interface with *R. alcyone* ACE2 is remodeled as compared to human ACE2 bound to SARS-CoV-2 or SARS-CoV-1, thereby explaining the shift in receptor species tropism.

### PRD-0038 RBD mutations enable human ACE2 utilization and expand receptor tropism to additional geographically-relevant bat species

To investigate how viral evolution could alter the PRD-0038 receptor species tropism for pandemic preparedness, we evaluated the impact of RBD mutations on utilization of several ACE2 orthologs. Using BLI, we observed that human ACE2-Fc did not bind to the wildtype PRD-0038 RBD (**Figure 2A**). However, we found that two amino acid residue substitutions promoted binding of human ACE2-Fc to the immobilized PRD-0038 K482Y/T487W RBD mutant (**Figure 2A**, SARS-CoV-2 numbering 493Y/498W))^29^, in line with their ability to enable BtKY72 S-mediated utilization of human ACE2^29,29^. We also found that human ACE2-Fc interacted with the T487W RBD, albeit weakly (**Figure 2A**). We validated these findings using deep mutational scanning (DMS) of the yeast-displayed PRD-0038 RBD showing that T487W is the only single amino acid mutation enabling detection of human ACE2-Fc binding (**Figures 2B and S3**). The geographically relevant *R. alcyone* and *R. landeri* ACE2 orthologs exhibited enhanced binding to the PRD-0038 T487W RBD relative to the wildtype RBD, with 1:1 binding affinity improvements corresponding to 2– and greater than one order of magnitude, respectively (**Figures 2C, 2D and Table S3**). These data concur with our DMS measurements showing that T487W mutation had the most marked positive effect on *R. landeri* ACE2 binding (**Figures 2B and S3**). Moreover, the PRD-0038 T487W RBD mutation enhanced binding to the *R. affinis* 787 allele as well as to the *R. sinicus* alleles recognized by the wildtype PRD-0038 RBD and enabled detectable binding to the *R. sinicus* WJ1 allele (**Figures S4A and S4B**). In contrast, the K482 mutation was deleterious for binding to *R. alcyone* and *R. landeri* ACE2 orthologs (**Figures 2E, 2F, S3, S4C, and S4D**). We observed that wildtype PRD-0038 S and all three mutants (K482Y, T487W and K482Y/T487W) promoted entry of VSV pseudotypes in HEK293T cells transiently transfected with *R. alcyone* ACE2 or with *R. landeri* ACE2 except for PRD-0038 K482Y S VSV that did not enter *R. landeri* ACE2-expressing cells (**Figures 2G, 2H, and S4E**), suggesting that binding avidity could overcome to some extent the observed differences in affinity. To broaden our understanding of PRD-0038 tropism we also examined cell entry promoted by transient transfection of ACE2 alleles from *R. ferrumequinum* bats, which are found in northern Africa, southern Europe, and southeast Asia. We observed that *R. ferrumequinum* ACE2 allele XM_033107295.1 promoted entry of wildtype and T487W PRD-0038 S VSV whereas *R. ferrumequinum* ACE2 allele FJ598617.1 enabled entry of T487W and even more so of K482Y/T487W PRD-0038 S VSV (**Figures 2I and 2J**). Finally, we observed entry of PRD-0038 T487W S and even more so of PRD-0038 K482Y/T487W S in HEK293T cells stably expressing human ACE2 (**Figure 2K**). These findings indicate that a single RBD residue mutation is sufficient for broadening the *Rhinolophus* bat ACE2 tropism and for enabling PRD-0038 S-mediated entry into cells expressing the human ACE2 receptor, highlighting the possible future zoonotic risk of this virus and related clade 3 sarbecoviruses.

**Figure 2.**
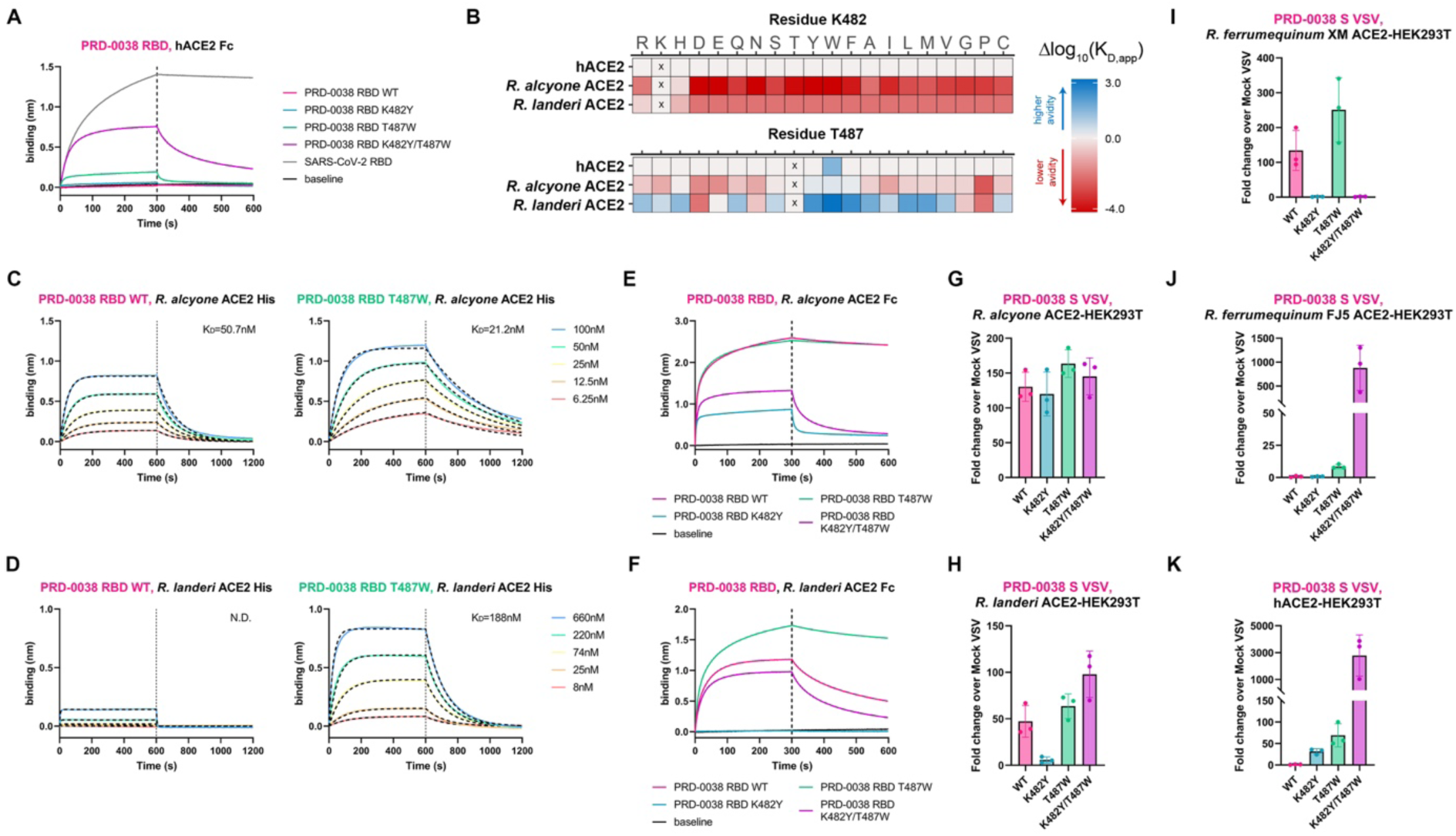
PRD-0038 RBD amino acid mutations broaden receptor tropism. (A) BLI binding analysis of 1 µM dimeric hACE2-Fc to biotinylated wildtype and mutant PRD-0038 RBDs immobilized on streptavidin biosensors. (B) DMS heatmaps of change in binding avidity to several ACE2-Fc orthologs caused by all possible mutations of the PRD-0038 RBD residues K482 and T487. An interactive version of the DMS data can be found at https://tstarrlab.github.io/SARSr-CoV-RBD_DMS/RBD-heatmaps_delta/. (C) BLI binding analysis of various concentrations of monomeric *R. alcyone* ACE2 to biotinylated wildtype (left) and T487W (right) PRD-0038 RBDs immobilized on streptavidin biosensors. (D) BLI binding analysis of various concentrations of monomeric *R. landeri* ACE2 to biotinylated wildtype (left) and T487W (right) PRD-0038 RBDs immobilized on streptavidin biosensors. (E) BLI binding analysis of 1 µM dimeric *R. alcyone* ACE2-Fc to biotinylated wildtype and mutant PRD-0038 RBDs immobilized on streptavidin biosensors. (F) BLI binding analysis of 1 µM dimeric *R. landeri* ACE2-Fc to biotinylated wildtype and mutant PRD-0038 RBDs immobilized on streptavidin biosensors. (G-K) Entry of VSV pseudotyped with wildtype and mutants PRD-0038 S into HEK293T cells transiently transfected with *R. alcyone* ACE2 (G), *R. landeri* ACE2 (H), *R. ferrumequinum* XM_033107295.1 (I, XM), *R. ferrumequinum* FJ598617.1 (J, FJ5), or stably expressing human ACE2 (K). See methods section and Figure S4C for VSV S pseudotype normalization details.

### Architecture of the PRD-0038 S trimer

To unveil the 3D organization of the clade 3 sarbecovirus infection machinery, we determined the structure of prefusion PRD-0038 S using single-particle cryoEM. After 3D classification and refinement, we obtained a 2.8 Å resolution reconstruction (**Figures 3A, 3B, 3C, S5, and Table S2**) of the trimer with the three RBDs in the closed conformation applying C3 symmetry, as we did not detect particle images corresponding to S trimers with open RBD conformations (**Figure S5**). We used local refinement to improve the resolution of the N-terminal domain (NTD) within the S trimer yielding a map at 2.9 Å resolution (**Figure S5**). The final model contains residues 18-1125 with chain breaks between residues 667-673 and 812-830.

**Figure 3.**
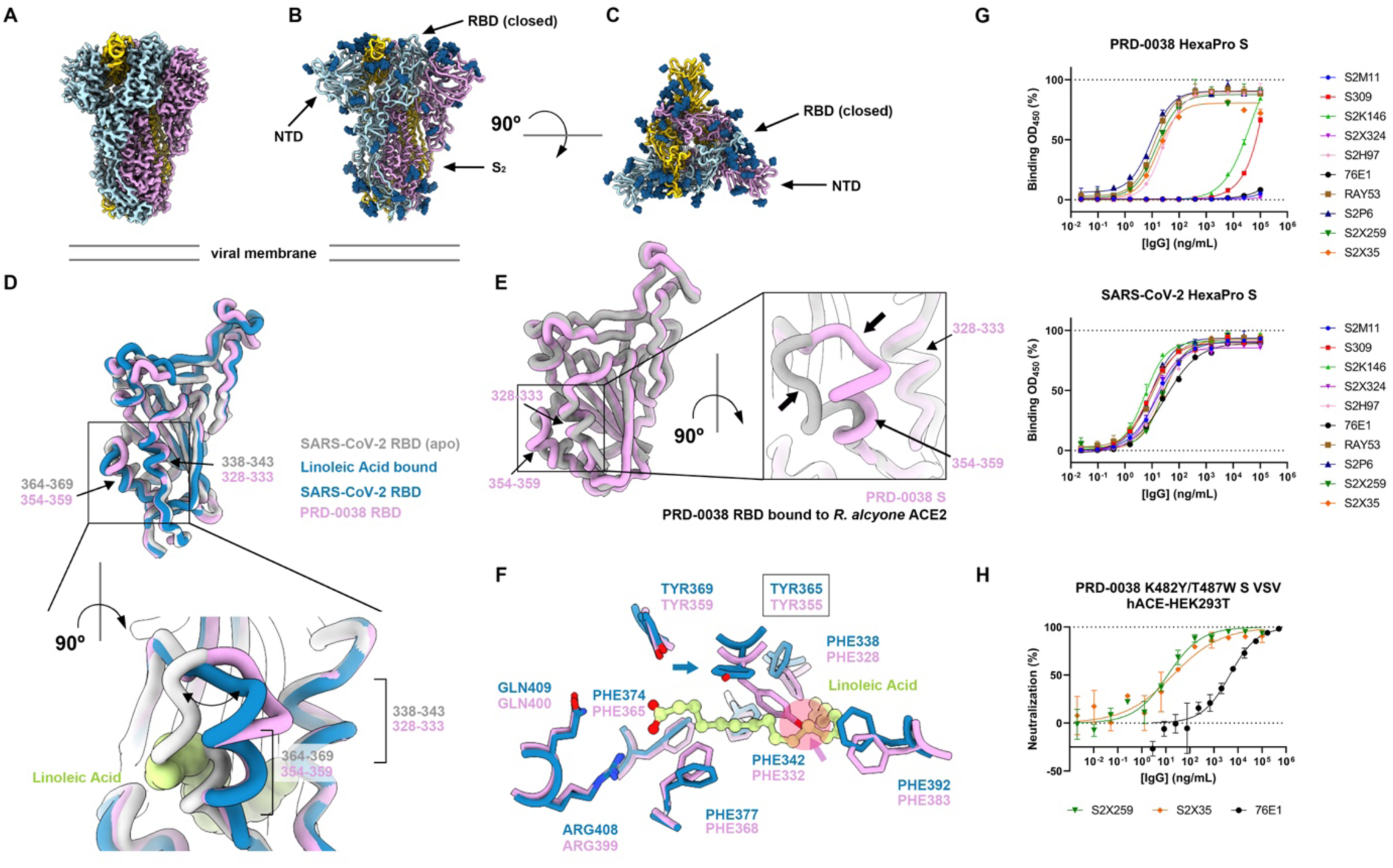
Architecture and antigenicity of the PRD-0038 S trimer. (A) Unsharpened cryo-EM map of the closed PRD-0038 S trimer at 2.8 Å resolution. (B and C) Ribbon diagram of the PRD-0038 S trimer atomic model viewed along (B, side) and normal (C, top) to the viral membrane. N-linked glycans are rendered as blue spheres. (D) Superimposition of the PRD-0038 S structure described here to the apo (gray, PDB 6VXX^5^) and linoleic acid-bound (blue, PDB 6ZB5^44^) SARS-CoV-2 S structures with a close-up view of the linoleic acid-binding pocket. N-linked glycans are omitted for clarity. (E) Superimposition of the PRD-0038 RBDs from the apo S structure (pink) and from the *R. aclyone* ACE2-bound RBD structure (gray). (F) Conservation of the SARS-CoV-2 linoleic acid-binding residues in PRD-0038 S. The PRD-0038 S Y355 side chain rotamer (Y365 in SARS-CoV-2 numbering) would sterically hinder linoleic acid binding (semi-transparent red circle denoted with an arrow). (G) Evaluation of binding of a panel of monoclonal antibodies to PRD-0038 S Hexapro S and SARS-CoV-2 Hexapro S measured by ELISA. (H) Monoclonal antibody-mediated neutralization of PRD-0038 K482Y/T487W S pseudotyped VSV entry into HEK293T cells stably expressing human ACE2.

The overall architecture of the PRD-0038 S trimer is similar to that of SARS-CoV-1 and SARS-CoV-2 and a PRD-0038 S protomer can be superimposed with r.m.s.d. values of 0.9 and 0.8 Å to SARS-CoV-1 S and SARS-CoV-2 S with which it shares 72% and 75% amino acid sequence identity, respectively (**Figure S6**). Glycosylation at RBD residue N360, which is present in most sarbecoviruses except SARS-CoV-2 (position N370) has been reported to favor the closed S conformation^39,40^, which could possibly explain the sole observation of closed PRD-0038 S trimers in our cryoEM dataset. However, glycosylation at this site is also present in SARS-CoV-1, which spontaneously adopts open RBD conformations^41–43^. The closed PRD-0038 S RBD conformation is most similar to the linoleic acid (LA)-bound form of SARS-CoV-2 S previously described^44^ (**Figures 3D and S6**). Indeed, the conformation of the RBD helix containing residues 354-359 (equivalent to SARS-CoV-2 residues 364-369) closely resembles that of the linoleic acid– bound SARS-CoV-2 S (**Figure 3D**) or that of F371-harboring SARS-CoV-2 Omicron variants^45–48^. This conformation, however, is distinct from that observed in the structure of the isolated PRD-0038 RBD bound to *R. alcyone* ACE2 described above (**Figure 3E**). No linoleic acid density is resolved in our cryoEM map although the RBD pocket that accommodates this ligand is conserved in the PRD-0038 S structure, including residues R399 and Q400 (equivalent to SARS-CoV-2 R408 and Q409 forming electrostatic interactions with the linoleic acid carboxylate). Instead, we found that the Y355 side chain (equivalent to SARS-CoV-2 Y365) partially obstructs the hydrophobic pocket which would have otherwise been occupied by the linoleic acid hydrocarbon tail (**Figure 3F**). The PRD-0038 S Y355 side chain rotamer resembles that of apo SARS-CoV-2 S (PDB 6VXX^5^) and apo SARS-CoV-1 S (PDB 5X58^43^), suggesting that this rotameric configuration is accessible to SARS-CoV-2 Y365 and likely changes to allow linoleic acid binding^44^.

### Antigenicity of the PRD-0038 S trimer

To define the antigenic landscape of clade 3 sarbecoviruses, we probed binding of a panel of monoclonal antibodies with broadly neutralizing activity against sarbecoviruses^21,22,45,49,50^ and α– and β– coronaviruses^51–55^ to prefusion-stabilized PRD-0038 S harboring the HexaPro mutations^56^ using an enzyme-linked immunosorbent assay (ELISA) (**Figure 3G**). We found that S2X259^22^ and S2X35^49^ (antigenic site II) as well as S2H97 (antigenic site V) cross-reacted with PRD-0038 S whereas S309^57^ (antigenic site IV) bound very weakly (**Figure 3G**). The markedly dampened S309 binding likely results from the E340_SARS-CoV-2_/Q330_PRD-0038_ escape substitution previously identified by deep mutational scanning of the yeast-displayed SARS-CoV-2 Wuhan-Hu-1 RBD^21^ (**Figure S7**). The S2X324^45^ antibody (antigenic site Ib), which neutralizes a broad panel of SARS-CoV-2 variant and resembles LY-COV1404^58^, recognizes the SARS-CoV-2 437-448 loop which is remodeled in the PRD-0038 RBD, explaining the lack of binding to PRD-0038 S (**Figure S7**). The S2K146^50^ antibody (antigenic site Ia), which contacts an epitope sharing several residues within the ACE2 binding site, bound very weakly to wildtype PRD-0038 S whereas S2M11^59^ (antigenic site Ia) did not bind at all, as a result of extensive RBM (receptor-binding motif) mutations (**Figure S7**). Nevertheless, we previously showed that S2K146 weakly neutralized VSV pseudotyped with BtKY72 S (clade 3) harboring the K482Y/T487W mutations (SARS-CoV-2 numbering 493Y/498W)^50^, underscoring the possible usefulness of this antibody if such mutations arose in related clade 3 sarbecoviruses.

The stem helix-targeting S2P6^53^ antibody and the RAY53^51^ antibody (recognizing the fusion machinery apex) cross-reacted as efficiently with PRD-0038 S as they did with SARS-CoV-2 S whereas the fusion peptide-directed 76E1^52^ antibody did not, possibly as a result of the F823_SARS-CoV-2_Y806_PRD-0038_ epitope mutation (which is shared with other clade 3 sarbecoviruses) along with the presence of the F800P_PRD-0038_ HexaPro stabilizing mutation^55^ (**Figures 3G and S7**).

Consistent with the ELISA data, we found that PRD-0038 K482Y/T487W S VSV pseudovirus was neutralized in a concentration-dependent manner by S2X259, S2X35 and more weakly by 76E1, the activity of the latter antibody is likely explained by the absence of the F800P_PRD-0038_ HexaPro stabilizing mutation (**Figures 3G and 3H**). Collectively, these data show that monoclonal antibodies targeting RBD antigenic sites or fusion machinery epitopes that are conserved across sarbecoviruses or α– and β-coronaviruses, respectively, retain neutralizing activity against PRD-0038 and are possible candidates for pandemic preparedness.

### Immunogenicity of the PRD-0038 S trimer

To better understand the immunogenicity of clade 3 sarbecoviruses and the impact of their possible inclusion in vaccine candidates, we immunized groups of six mice with three 1 µg doses of either PRD-0038 S or SARS-CoV-2 S (**Figure 4A**), both stabilized in the prefusion conformation using the HexaPro mutations^56^. Serum neutralizing activity was analyzed two weeks post dose 3 using VSV particles pseudotyped with clade 1a (SARS-CoV-1), clade 1b (SARS-CoV-2/G614, BA.2, BA.5) or clade 3 (PRD-0038 and Khosta1) S glycoproteins. SARS-CoV-2 S-immunized mice had potent serum neutralizing activity against SARS-CoV-2/G614 VSV S (vaccine-matched), which was reduced against BA.2 S VSV and even more so against BA.5 S VSV (**Figure 4B and S8**). RaTG13 S VSV (vaccine-mismatched), however, was neutralized with almost comparable potency to that against SARS-CoV-2/G614 S VSV whereas no neutralization of PRD-0038 S VSV and Khosta1 S VSV clade 3 sarbecoviruses (vaccine-mismatched) was detected except for one animal. PRD-0038 S-immunized mice had potent serum neutralizing activity against PRD-0038 S VSV (vaccine-matched) and Khosta1 S VSV (vaccine-mismatched) whereas we could not detect any neutralization of clade 1a and 1b pseudoviruses tested besides weak RaTG13 inhibition (**Figure 4B and S8**). Although none of the sera could block SARS-CoV-1 S-mediated entry into target cells in standard experimental conditions, we observed weak SARS-CoV-1 S VSV neutralization with greater dilution of the pseudovirus stock when using PRD-0038 S-but not SARS-CoV-2 S-elicited sera (**Figure S9**). These results suggest that this clade 3 S trimer immunogen induced more broadly reactive antibody responses than SARS-CoV-2 S as judged by cross-clade reactivity.

**Figure 4.**
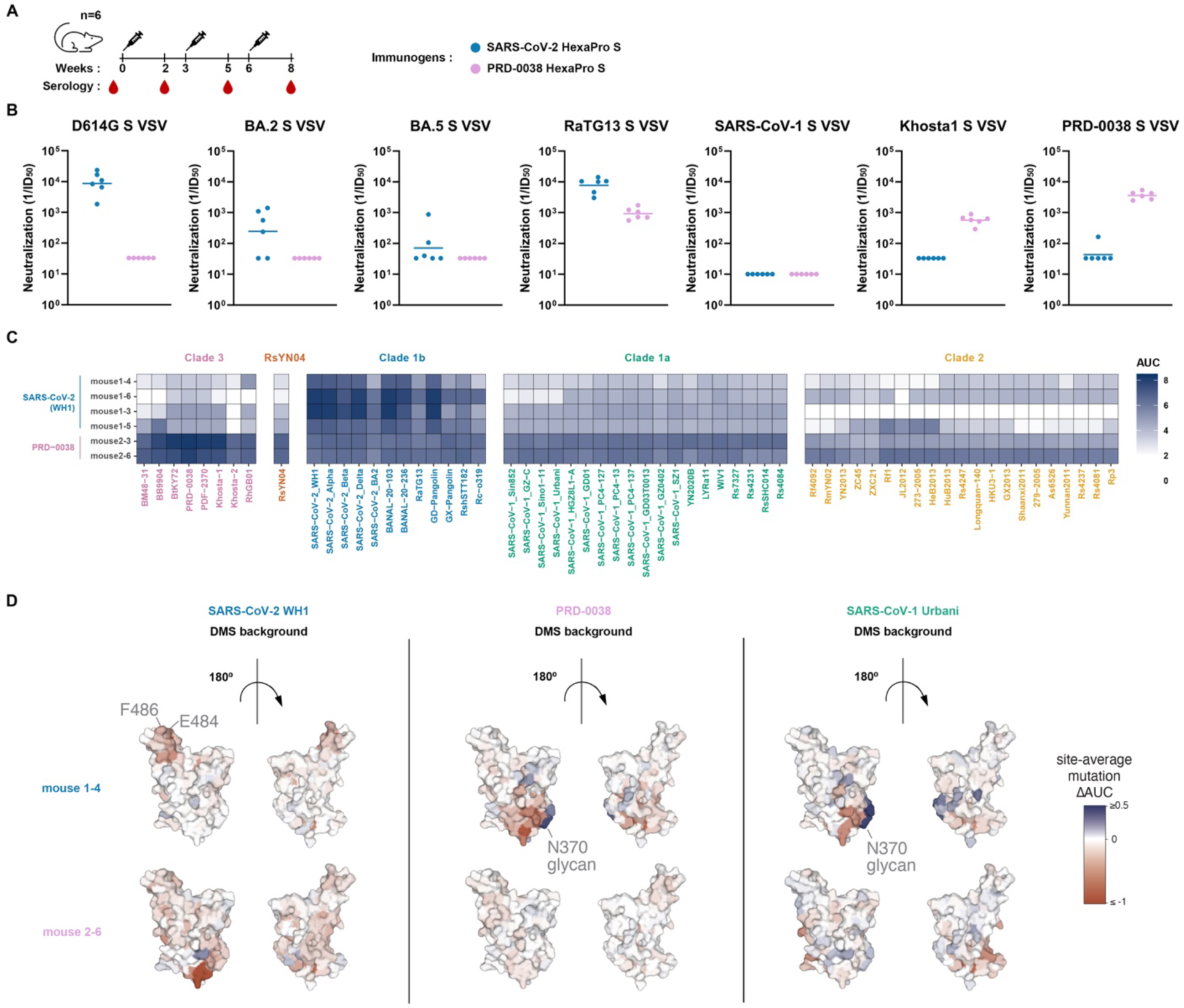
A clade 3 sarbecovirus S trimer elicits broadly reactive antibody responses. (A) Vaccination schedule for the mouse immunogenicity study. (B) Serum neutralization of VSV pseudotyped with various sarbecovirus S glycoproteins. Bar represents the geometric mean of each group. (C) Sarbecovirus breadth of serum binding to a pan-sarbecovirus library of yeast-displayed RBDs using a high-throughput FACS-seq assay. (D) Epitope targeting of serum antibodies. For two representative sera (mouse 1-4, vaccinated with SARS-CoV-2 HexaPro S and mouse 2-6, vaccinated with PRD-0038 HexaPro S), we determined the dominant antibody epitopes via DMS using vaccine-matched and mismatchedRBD backgrounds. The average effect of mutations at each site are mapped to the SARS-CoV-2 structure, where blue and red indicate positions where mutations increase or decrease serum binding, respectively. See Figure S11 for all DMS profiles.

We subsequently analyzed binding of selected vaccine-elicited sera to a panel of yeast-displayed RBDs spanning the known sarbecovirus phylogenetic diversity using two mice immunized with PRD-0038 S and four mice immunized with SARS-CoV-2 S. Mice were chosen based on neutralization potency against BA.2, BA.5, or SARS-CoV-1 S VSV. We selected two mice with greatest (mouse 1-3, 1-5) and two mice with weakest (mouse 1-4, 1-6) serum neutralizing activity from the group immunized with SARS-CoV-2 HexaPro S as well as two mice with the greatest SARS-CoV-1 cross-neutralization (mouse 2-3, 2-6) from the group immunized with PRD-0038 HexaPro S (**Figure S8**). In line with the serum neutralization data, inclusion of an antigen in the vaccine formulation was associated with strong cross-reactivity with vaccine-matched and related antigens within the same clade among the sera analyzed (**Figure 4C**). Furthermore, PRD-0038 S immunization elicited greater titers of antibodies cross-reacting with clade 2 and clade 1a RBDs, as well as the RsYN04 RBD that branches independently of the four previously known clades^60^, likely explaining the weak but detectable SARS-CoV-1 neutralization (**Figures 4B, 4C, and S9**). These data indicate that inclusion of a clade 3 antigen in a vaccine formulation could not only elicit clade 3 serum neutralizing activity but also enhance cross-reactive (and weakly neutralizing) antibody responses against vaccine-mismatched antigens from distinct clades, which could participate in protection through direct neutralization and Fc-mediated effector functions^46,61–64^.

To understand the molecular basis for variation in breadth of serum cross-reactivity across mice and vaccine regimen, we mapped the dominant epitope specificities in these six sera using yeast-displayed DMS libraries in the vaccine-matched SARS-CoV-2 Wuhan-Hu-1 or PRD-0038 RBDs and in the vaccine-mismatched SARS-CoV-2 Omicron BA.2 and SARS-CoV-1 Urbani RBDs (**Figures 4D and S10**). SARS-CoV-2 S-elicited polyclonal serum antibodies predominantly targeted the SARS-CoV-2 RBD region comprising residues 484-490, consistent with the strong antigenic pressure on this position that drove early variant evolution at residue E484 during the COVID-19 pandemic^65–67^. These residues are highly variable among sarbecoviruses, consistent with the weaker cross-reactive breadth seen in these sera. In contrast, PRD-0038 S-elicited polyclonal serum antibodies showed no dominant targeting of specific PRD-0038 RBD antigenic sites, which could indicate a more balanced binding antibody response that would be less susceptible to single amino acid mutations. Epitope mapping to vaccine-mismatched RBDs revealed that PRD-0038 and SARS-CoV-2 S-elicited sera contained subdominant antibodies targeting antigenic sites II, IV and V^21,49^, which are typically recognized by broadly reactive and neutralizing antibodies^21,22,57,68,69^. However, DMS using vaccine-mismatched RBD strains also reveals the presence of antibodies in SARS-CoV-2 S-elicited sera that target suboptimal sites for sarbecovirus breadth. For example, we observed strong antibody responses to the N370 glycan hole (resulting from mutations of this glycosylation sequon in our DMS experiments) for two of the four SARS-CoV-2-immunized mice analyzed. Antibody responses targeting this site are elicited as a result of the absence of this oligosaccharide in the SARS-CoV-2 RBD immunogen but its presence in all other sarbecovirus RBDs. Restoration of the N370 glycan in SARS-CoV-2 vaccines could therefore possibly limit these off-target responses by reducing elicitation of antibodies to this strain-specific epitope. In conclusion, the DMS data indicate that PRD-0038 S-elicited serum antibodies analyzed here target a broader spectrum of antigenic sites present on vaccine-matched and mismatched RBDs than SARS-CoV-2 S-elicited antibodies, providing a molecular basis for enhanced cross-reactivity (**Figures 4D and S10**).

## DISCUSSION

Coronavirus S glycoproteins are evolutionary hotspots and can acquire amino acid substitutions, insertions, deletions or even recombine distinct domains^70^. These mutational changes can alter host receptor tropism^71^, binding affinity^38,72,73^, entry route ^46,74,75^ and immune evasion^36,37,72,76–78^. Most mutations occur within the RBDs which engage the host receptor and account for most neutralizing activity against vaccine/infection-matched and mismatched sarbecoviruses^49,79–81^.

Spillover is a complex process involving multiple factors such as receptor recognition, proteolytic S activation, immune antagonism and contact opportunity. To examine potential spillover pathways of clade 3 sarbecoviruses, we evaluated binding of PRD-0038 RBD mutants to a panel of *Rhinolophus* bat ACE2 orthologs. Although human ACE2 cannot serve as entry receptor for wildtype PRD-0038 S (or the closely related BtKY72 S^29^), introduction of a single amino acid RBD mutation (T487W) enabled binding and S-mediated entry into cells expressing human ACE2. Moreover, this point mutation broadened host receptor tropism by enabling utilization of the geographically-relevant *R. landeri* and *R. ferrumequinum* ACE2 receptors without compromising binding to *R. alcyone*, *R. sinicus* and *R. affinis* ACE2 alleles. Although the T487W RBD mutation requires 3 nucleotide substitutions, these findings point to a possible spillover pathway in which a single amino acid change expands host receptor tropism markedly. Indeed, acquisition of *R. landeri*, *R. ferrumequinum* (allele FJ598617.1) and human ACE2 tropism would allow PRD-0038 (and likely BtKY72 and related clade 3 viruses) to expand the geographic range of host reservoirs that can be infected and in turn the likelihood of zoonotic transmission. Broadly neutralizing antibodies with activity against sarbecovirus (RBD-directed) and beyond (fusion machinery-directed) inhibited PRD-0038 S-mediated entry into cells and could be stockpiled as possible countermeasures for pandemic preparedness.

We observed that three immunizations with a clade 1b or with a clade 3 sarbecovirus S trimer predominantly elicited vaccine-matched serum neutralizing antibody responses. We note that our neutralization data of clade 1b pseudoviruses underscore the distinction between antigenic and genetic distance: although RaTG13 harbors a greater number of RBD mutations than Omicron BA.2 or BA.5, neutralizing activity was higher against RaTG13 than against these SARS-CoV-2 variants which accumulated mutations to erode neutralizing antibody titers^52,82^. Compared to SARS-CoV-2 S, we found that PRD-0038 S-elicited polyclonal serum antibodies were more broadly reactive with vaccine-mismatched antigens, including clades 1a and 2 RBDs, which likely account for the weak but detectable cross-neutralization of SARS-CoV-1 with sera from mice immunized with PRD-0038 S but not with SARS-CoV-2 S. As non– or weakly neutralizing monoclonal and polyclonal antibodies have been shown to participate in protection against SARS-CoV-2 challenge in small animal models through Fc-mediated effector functions^46,61–64^, our findings motivate the inclusion of clade 3 and other divergent RBD antigens in updated sarbecovirus vaccine formulations^83–86^. This would allow elicitation of potent clade 3 neutralizing antibodies and cross-clades binding (and possibly neutralizing) antibodies with maximal breadth to achieve optimal protection against continuously evolving SARS-CoV-2 variants and sarbecoviruses found in wildlife.

## Limitations of the study

There is currently no known animal challenge model for clade 3 sarbecoviruses and we were therefore not able to evaluate the contribution of the broadly reactive antibody responses elicited upon PRD-0038 S vaccination to in vivo protection.

## Acknowledgements

This study was supported by the National Institute of Allergy and Infectious Diseases (F31AI174573-01 to S.K.Z., P01AI167966 to N.P.K, T.N.S. and D.V., DP1AI158186 and 75N93022C00036 to D.V., K99AI166250 to T.N.S.), the National Institute of General Medical Sciences (T32GM008268-32 to S.K.Z.), a Pew Biomedical Scholars Award (D.V.), an Investigators in the Pathogenesis of Infectious Disease Awards from the Burroughs Wellcome Fund (D.V.), a Dale F. Frey Award for Breakthrough Scientists from the Damon Runyon Cancer Research Foundation (T.N.S.), the University of Washington Arnold and Mabel Beckman cryoEM center and the National Institute of Health grant S10OD032290 (to D.V.). D.V. is an Investigator of the Howard Hughes Medical Institute and the Hans Neurath Endowed Chair in Biochemistry at the University of Washington. We thank the High-Throughput Genomics Shared Resource at the University of Utah Huntsman Cancer Institute supported by NCI/NIH (P30CA042014), the Flow Cytometry Core Facility at the University of Utah Health Sciences Campus supported by NIH (S10OD026959 and 5P30CA042014-24), the University of Utah Center for High Performance Computing, supported by NIH (1S10OD021644-01A1), and the Fred Hutchinson Cancer Center Genomics core facility for experimental support.

## Author Contributions

J.L, S.K.Z., T.S., and D.V. designed the experiments; J.L, S.K.Z. and C.S. recombinantly expressed and purified glycoproteins. J.L and S.K.Z. performed binding assays. J.L carried out pseudovirus entry assays. J.L., S.K.Z, J.Q. and Y.J.P. carried out cryoEM specimen preparation, data collection and processing of PRD-0038 S. J.L., S.K.Z, and Y.J.P. carried out cryoEM specimen preparation, data collection and processing of the PRD-0038 RBD bound to ACE2. J.L., Y.J.P. and D.V built and refined atomic models. E.M.L. and C.T. performed mouse immunizations and blood draws. A.L.T. and T.N.S. carried out DMS experiments. D.C. contributed unique reagents. J.L. and D.V. analyzed the data and wrote the manuscript with input from all authors; N.P.K.,T.N.S. and D.V. supervised the project.

## Figure legends

**Figure S1.**
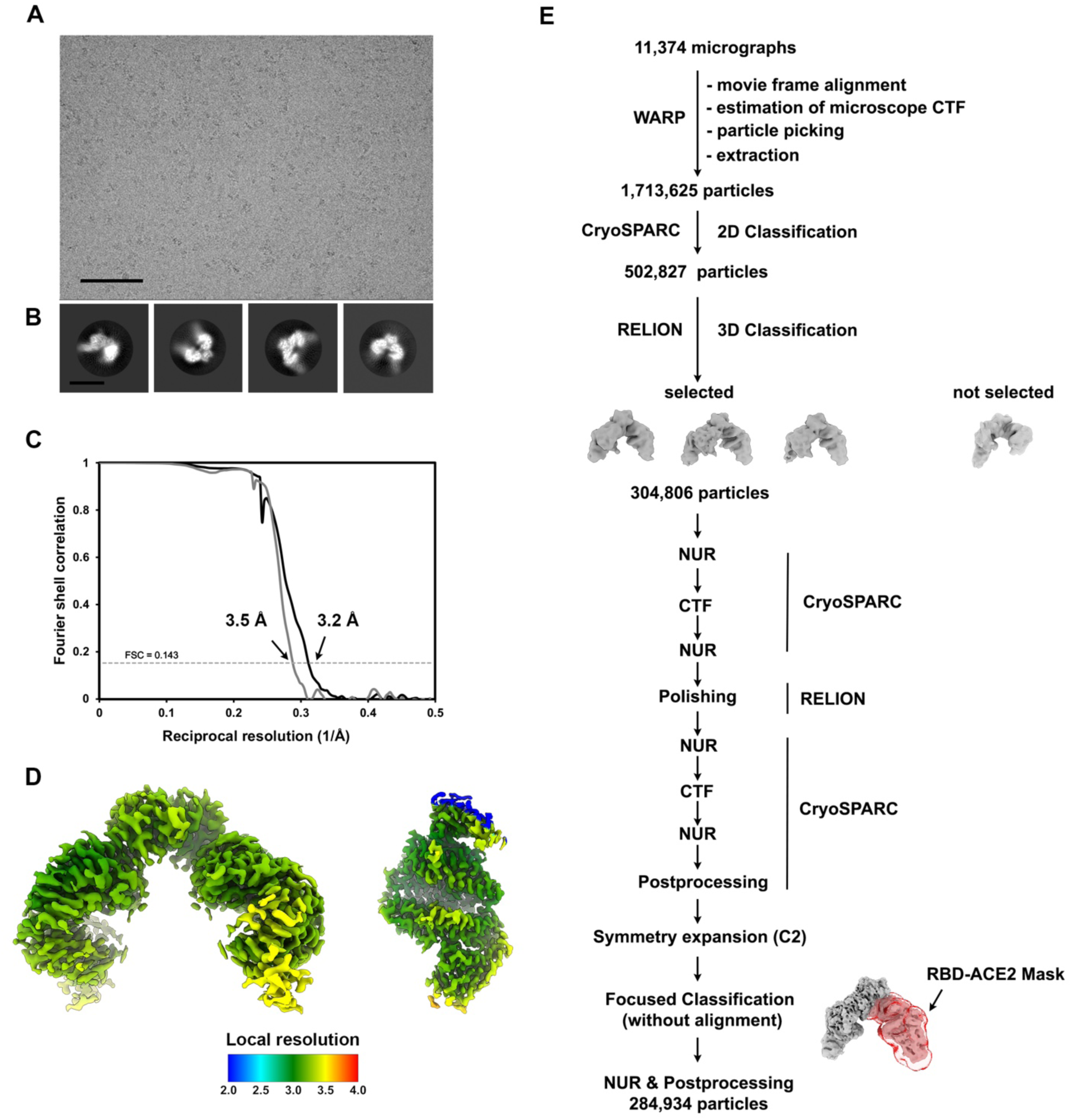
CryoEM data collection and refinement of the dimeric *R. alcyone* ACE2-bound PRD-0038 RBD complex. (A and B) Representative electron micrograph (A) and 2D class averages (B) of –the dimeric *R. alcyone* ACE2-bound PRD-0038 RBD embedded in vitreous ice. The scale bar represents 100 nm (A) or 100 Å (B). (C) Gold-standard Fourier shell correlation curves for the final cryoEM reconstructions of the dimeric –ACE2/RBD complex (solid gray line) and locally refined ACE2/RBD (solid black line) shown in (D). The 0.143 cutoff is indicated with a gray dashed line. (D) Local resolution map calculated using CryoSPARC and plotted onto the sharpened cryoEM map. (E) Data processing flowchart. CTF: contrast transfer function; NUR: non-uniform refinement.

**Figure S2.**
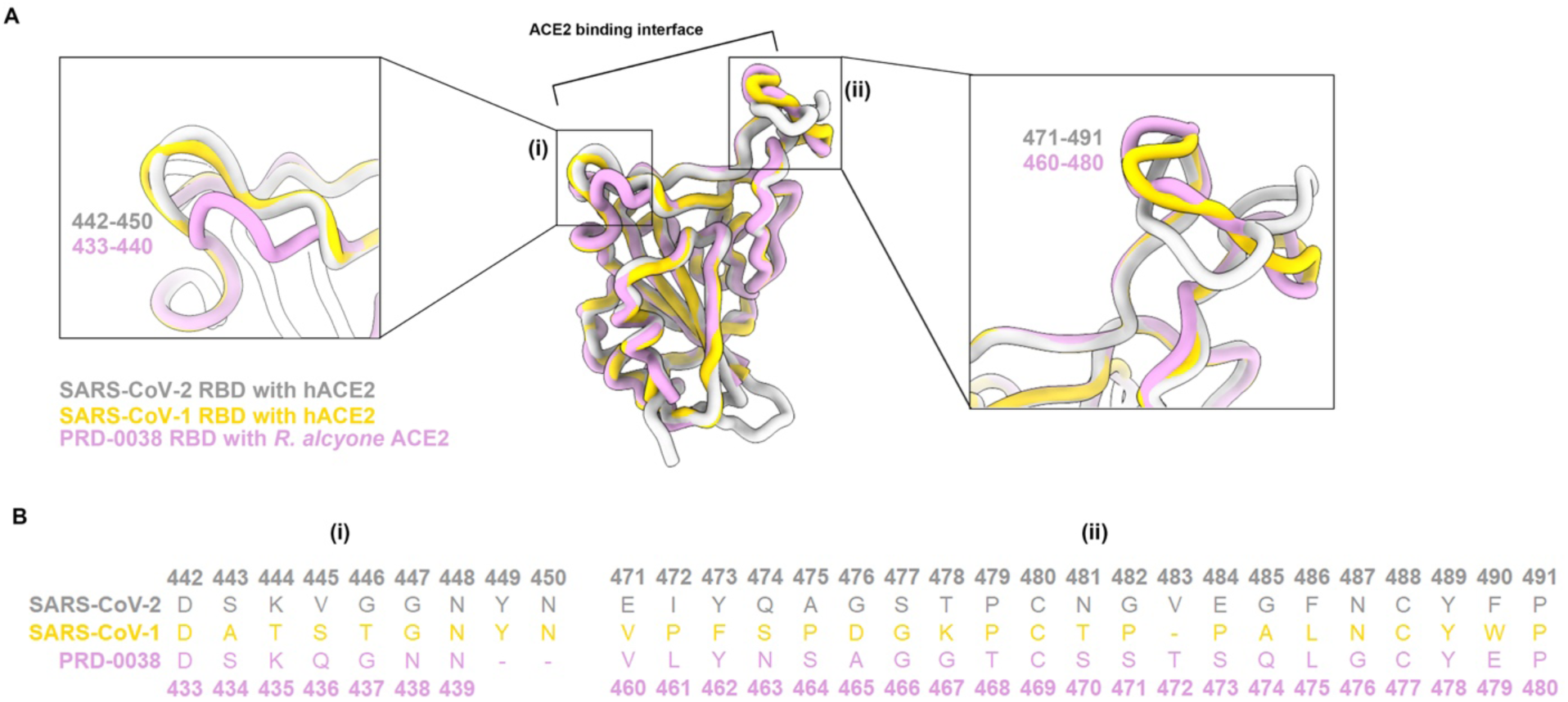
Structural distinctions between the PRD-0038, SARS-CoV-2 and SARS-CoV-1 RBDs near the ACE2-binding interface (RBM). (A) Ribbon diagrams showing a superimposition of the *R. alcyone* ACE2-bound PRD-0038 RBD (pink) superimposed to the human ACE2-bound SARS-CoV-2 RBD (gray, PDB 6M0J^33^) and SARS-CoV-1 RBD (gold, PDB 2AJF^34^) structures (ACE2s not shown for clarity). Insets: close-up views of two RBM regions. (B) Amino acid sequence alignment of the SARS-CoV-2, SARS-CoV-1, and PRD-0038 RBD regions highlighted in the insets shown in panel (A). – indicate deletions.

**Figure S3.**
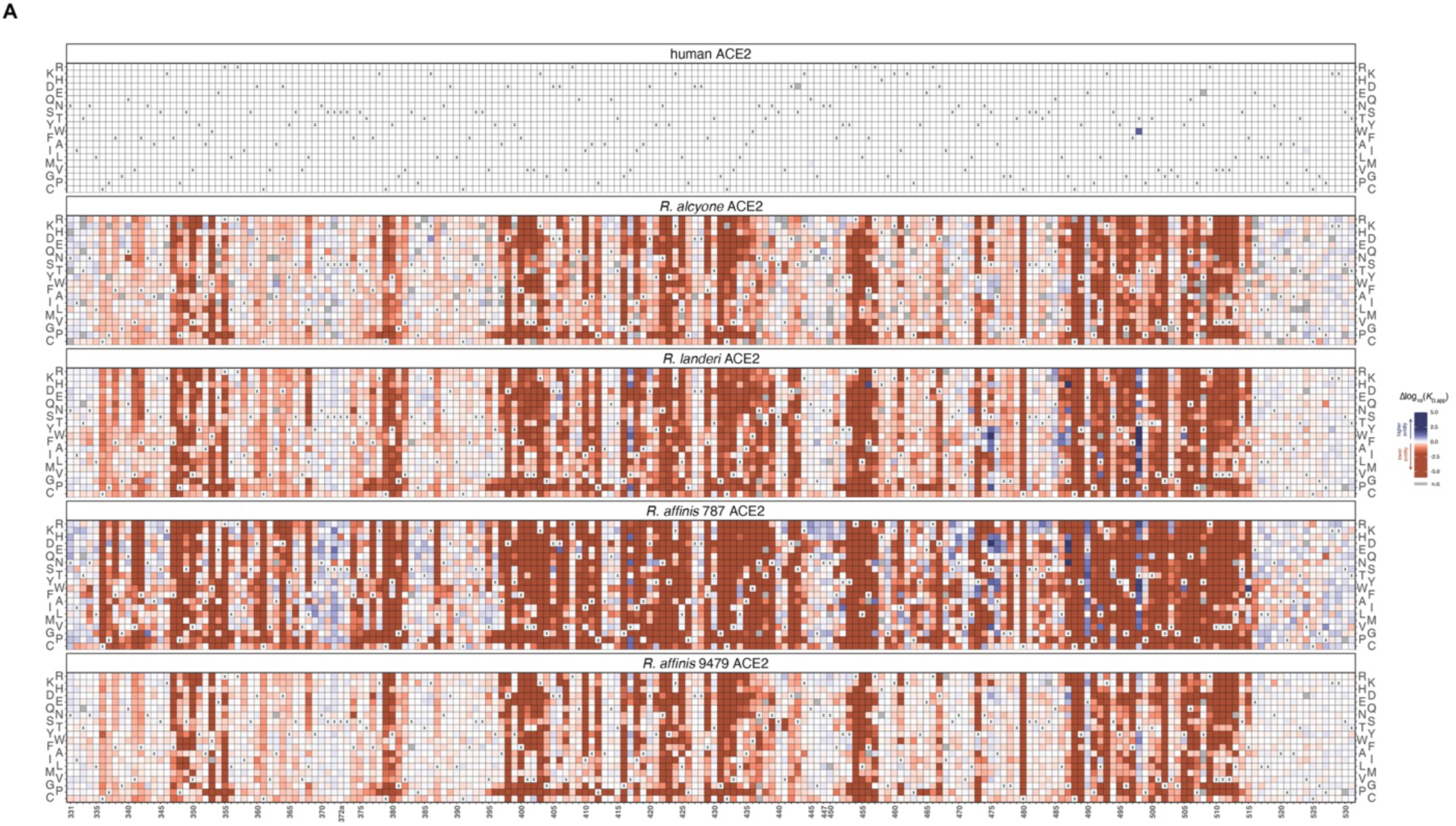
Heatmaps of change in ACE2-binding avidity resulting from RBD mutations determined by DMS. (A) Change in ACE2-binding avidity for hACE2, *R. alcyone* ACE2, and *R. landeri* ACE2 (delta-log_10_K_D_,apparent). Residue numbering corresponds to SARS-CoV-2.

**Figure S4.**
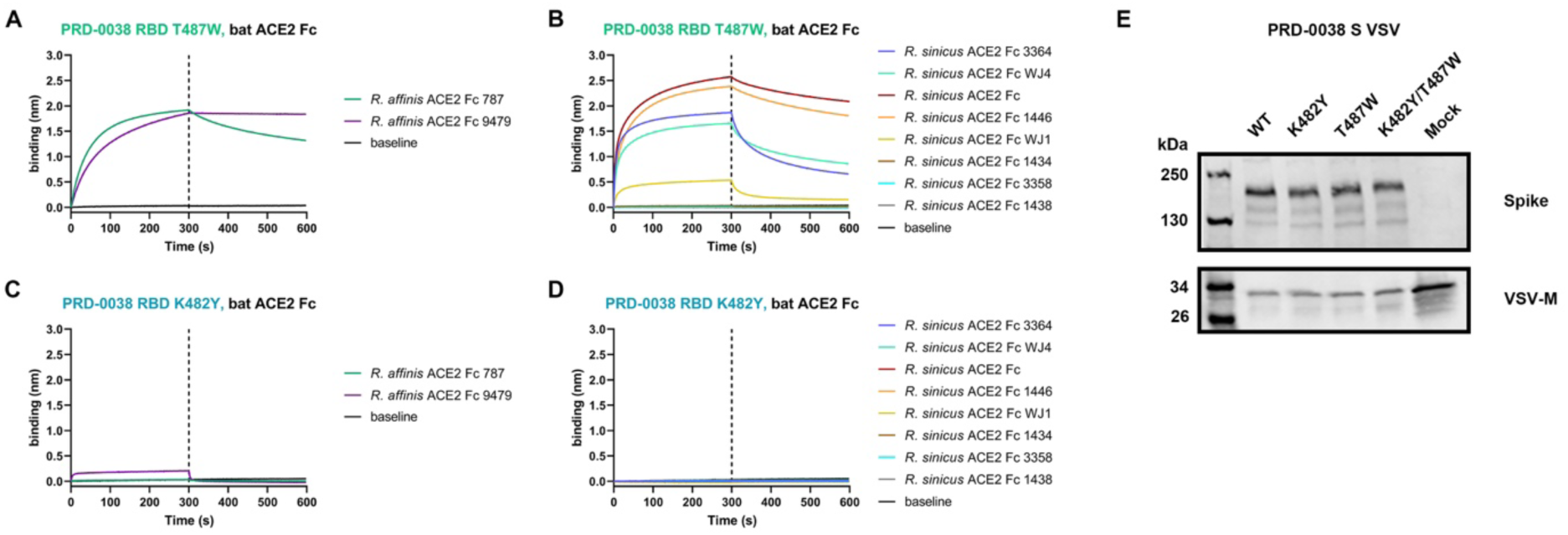
Binding data and Western blot analysis. (A and B) BLI binding analysis of 1 µM dimeric *R. affinis* (A) and *R. sinicus* (B) ACE2-Fc alleles to the biotinylated T487W PRD-0038 RBD immobilized on streptavidin biosensors. (C and D) BLI binding analysis of 1 µM dimeric *R. affinis* (C) and *R. sinicus* (D) ACE2-Fc alleles to the biotinylated K482Y PRD-0038 RBD immobilized on streptavidin biosensors. (E) Representative Western Blot of wildtype (WT) and mutant PRD-0038 S VSV pseudoviruses normalized based on the amount of incorporated S and VSV-M. Anti-VSV-M Antibody (Kerafast) and Monoclonal ANTI-FLAG® M2 antibody produced in mouse (Sigma) were used as the primary antibody against VSV backbone and S, respectively. Alexa Fluor® 680 AffiniPure Goat Anti-Mouse IgG (Jackson ImmunoResearch) was used as the secondary antibody.

**Figure S5.**
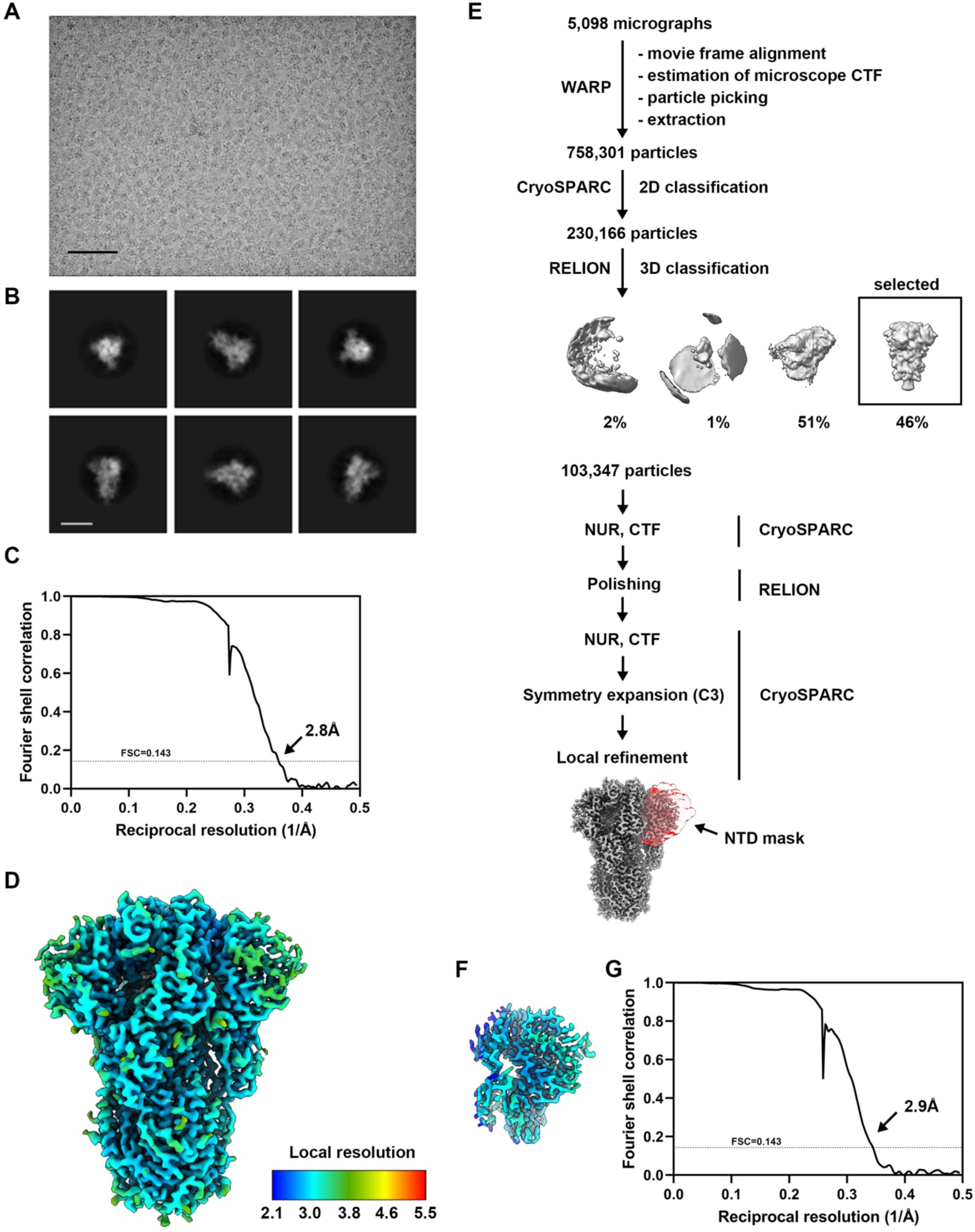
CryoEM data collection and refinement of PRD-0038 S. (A and B) Representative electron micrograph (A) and 2D class averages (B) of PRD-0038 PentaPro S embedded in vitreous ice. The scale bar represents 100 nm (A) or 100Å (B). (C) Gold-standard Fourier shell correlation curve for the cryoEM reconstruction. The 0.143 cutoff is indicated with a gray dashed line. (D) 3D reconstruction of PRD-0038 PentaPro S colored by local resolution as determined using cryoSPARC. (E) Data processing flowchart. CTF: contrast transfer function; NUR: non-uniform refinement. (F) 3D reconstruction obtained by local refinement of the PRD-0038 S NTD colored by local resolution as determined using cryoSPARC. (G) Gold-standard Fourier shell correlation curve. The 0.143 cutoff is indicated with a gray dashed line.

**Figure S6.**
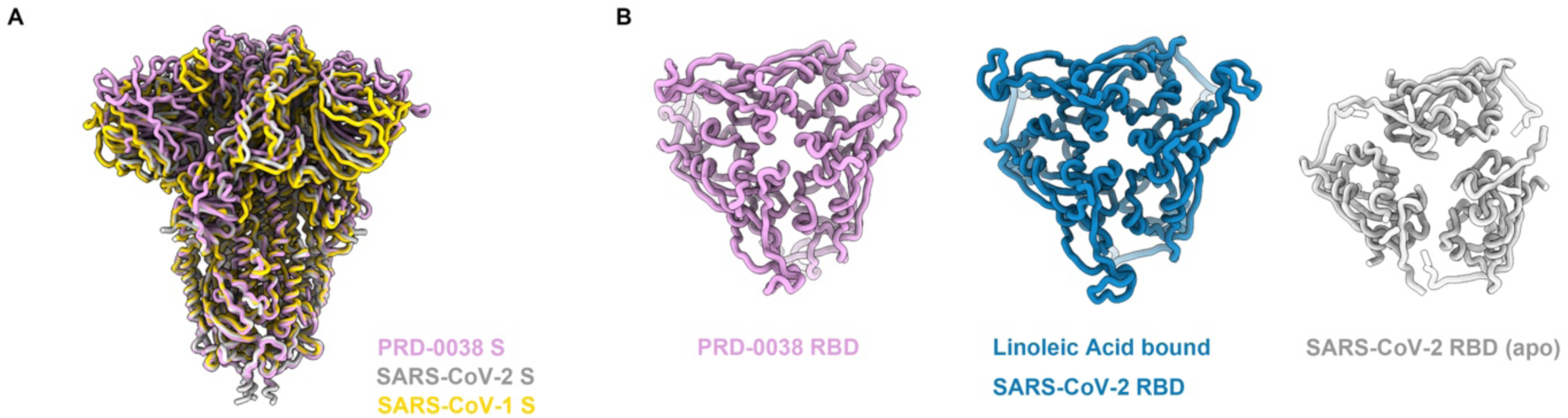
Structural resemblance of PRD-0038 S to SARS-CoV-2 and SARS-CoV-1. (A) Ribbon diagram of the PRD-0038 S trimer superimposed to SARS-CoV-2 S (PDB 6VXX^5^) and SARS-CoV-1 S (PDB 5X5B^43^). (B) Contact among RBDs within S trimers viewed from the apex along the 3-fold molecular axis of PRD-0038 S (pink), linoleic acid-bound SARS-CoV-2 S (blue, PDB 6ZB5^44^), and apo SARS-CoV-2 S (gray, PDB 6VXX^5^).

**Figure S7.**
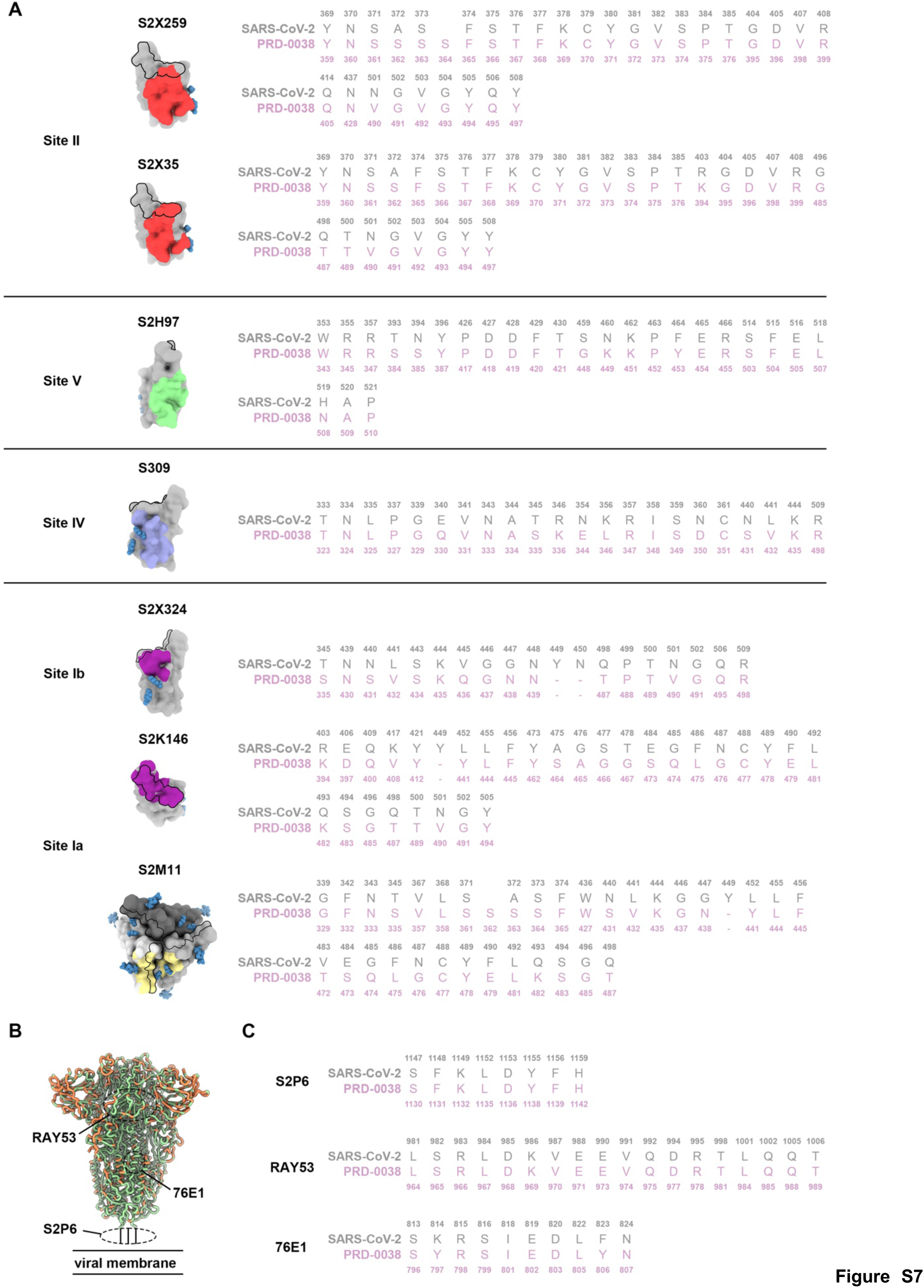
Conservation analysis of epitopes targeted by monoclonal antibodies between PRD-0038 S and SARS-CoV-2 S. (A) Epitope mapped onto PRD-0038 RBD structure with sequence alignment of key residues at the interface. The PRD-0038 RBD is shown in gray (three RBDs are shown with distinct shades of gray for S2M11 which recognizes a quaternary epitope) and N-linked glycans are rendered as blue spheres. SARS-CoV-2 residue numbering is shown in gray and PRD-0038 residue numbering is shown in pink. (B) PRD-0038 S sequence conservation with SARS-CoV-2 S at key S_2_ fusion machinery epitopes (indicated with dashed lines). Residues are colored according to sequence identity (orange: not conserved, green: conserved). (C) Sequence alignment of corresponding S_2_ epitopes. SARS-CoV-2 residue numbering is shown in gray and PRD-0038 residue numbering is shown in pink. For (A) and (C) the RBM is depicted with a black outline and the monoclonal antibody epitopes are colored according to their antigenic sites (I, purple; II, red; IV, violet; V, green).

**Figure S8.**
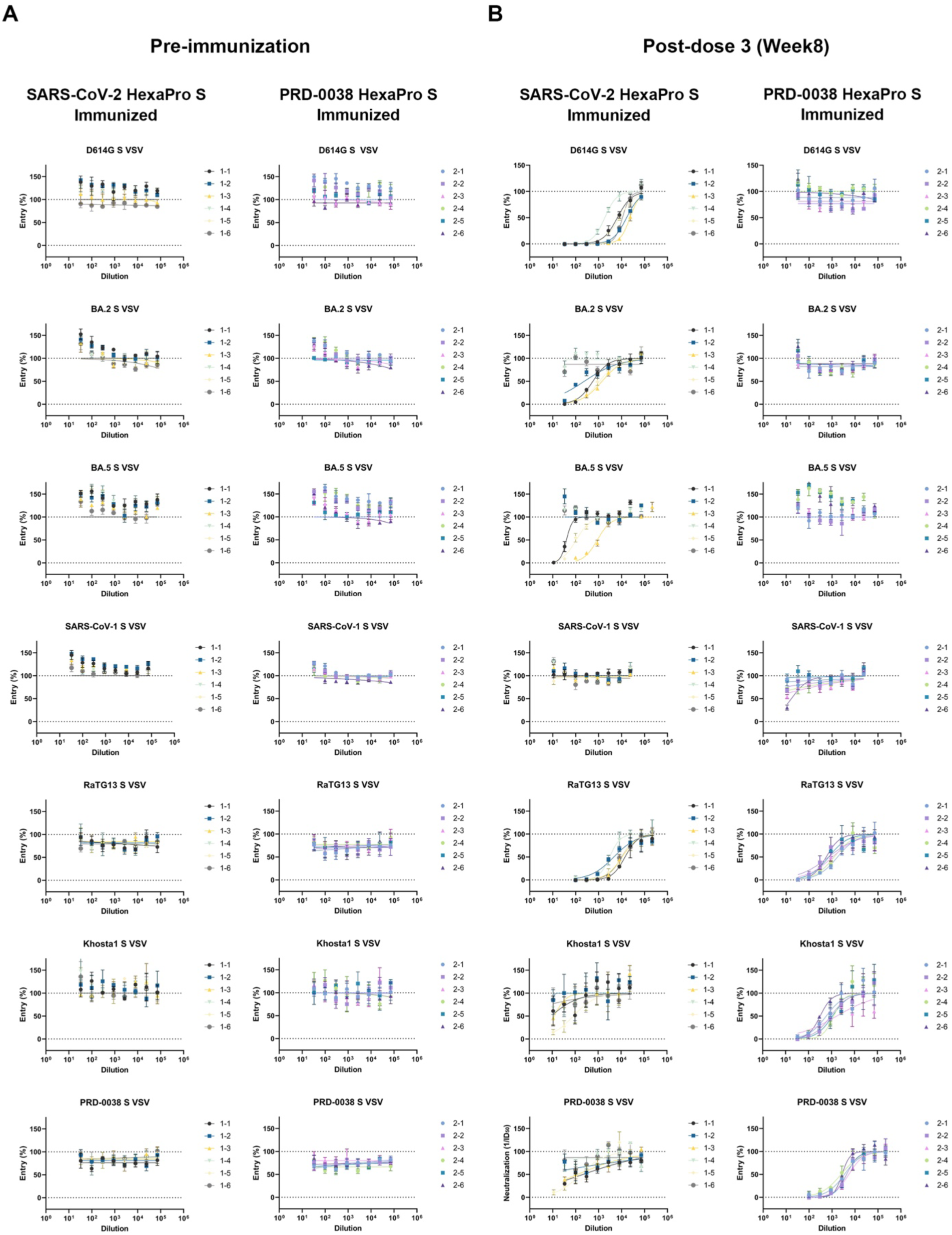
Dose-response curves of mouse serum neutralization before (A) and after (B) three immunizations with SARS-CoV-2 HexaPro S or PRD-0038 HexaPro S. Neutralization of the indicated sarbecovirus S VSV pseudotypes was assessed for each animal serum two weeks post dose 3 (week 8), as indicated by the color key.

**Figure S9.**
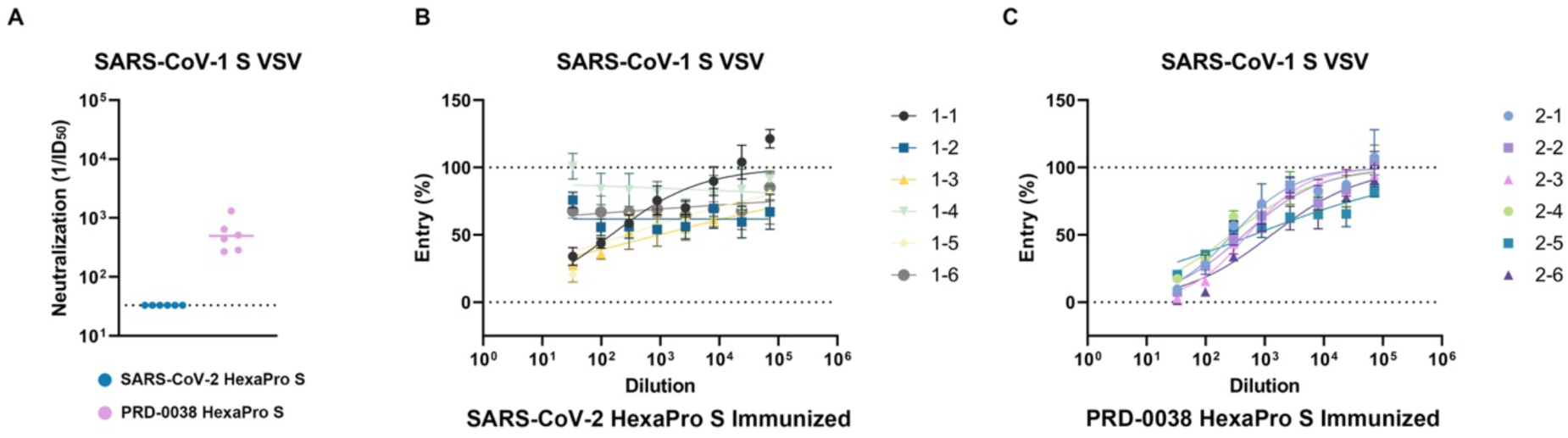
Neutralization of SARS-CoV-1 S VSV by vaccine-elicited mouse sera using a highly diluted pseudovirus input. (A) Reciprocal ID_50_ value. (B and C) Dose-response curves for neutralization of SARS-CoV-1 S VSV by SARS-CoV-2 HexaPro S-elicited sera (B) and PRD-0038 HexaPro S-elicited sera (C).

**Figure S10.**
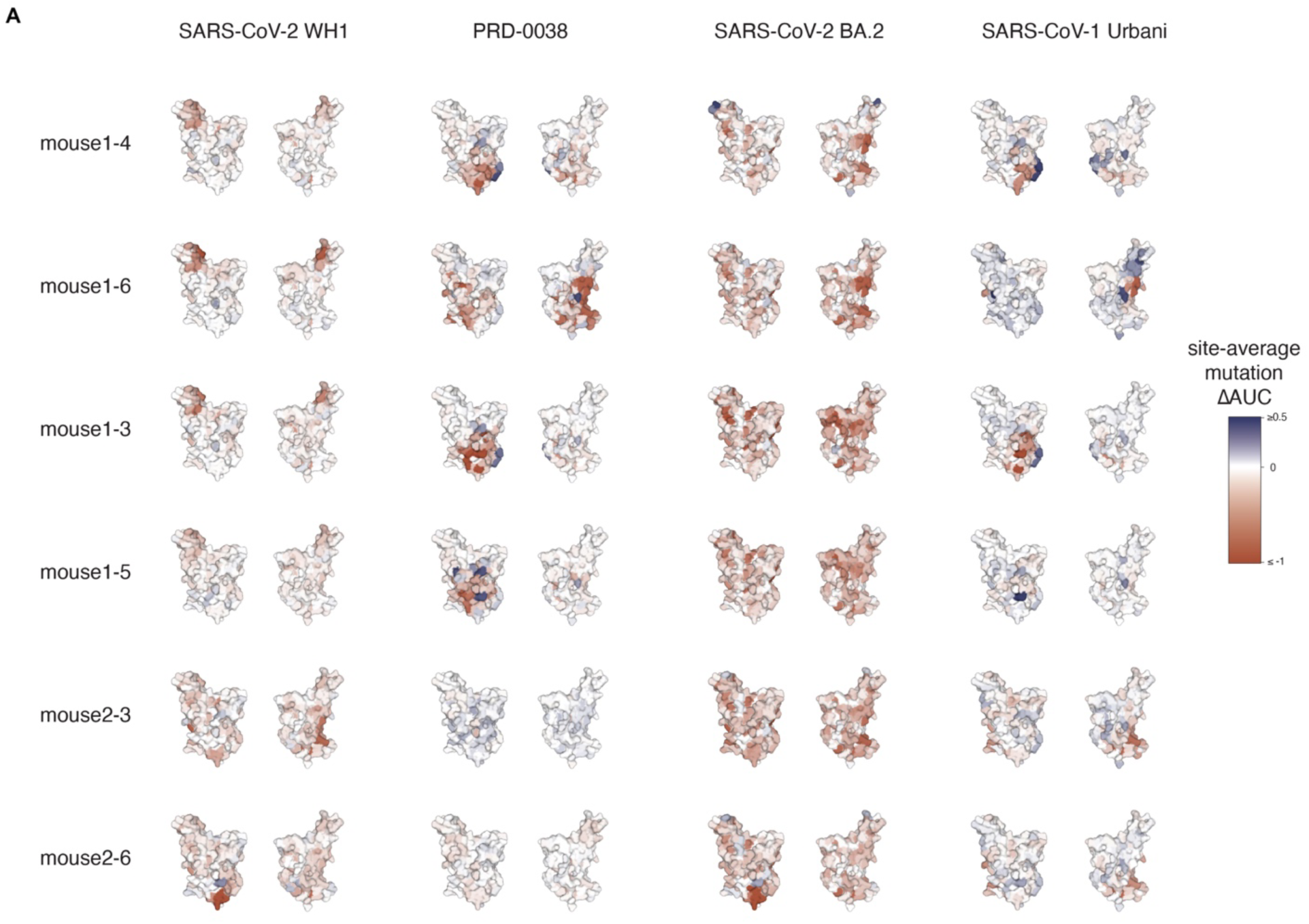
DMS epitope mapping of serum antibodies. (A) Evaluation of epitopes targeted by serum antibodies elicited by SARS-CoV-2 HexaPro S (mouse 1-3, 1-4, 1-5, and 1-6) or PRD-0038 HexaPro S (mouse 2-3 and 2-6) vaccination using yeast-displayed DMS of vaccine-matched and mismatched RBDs (indicated above each column). The average effect of mutations at each site are mapped to the SARS-CoV-2 structure, where blue and red indicate positions where mutations increase or decrease serum binding, respectively.

**Table S1.**
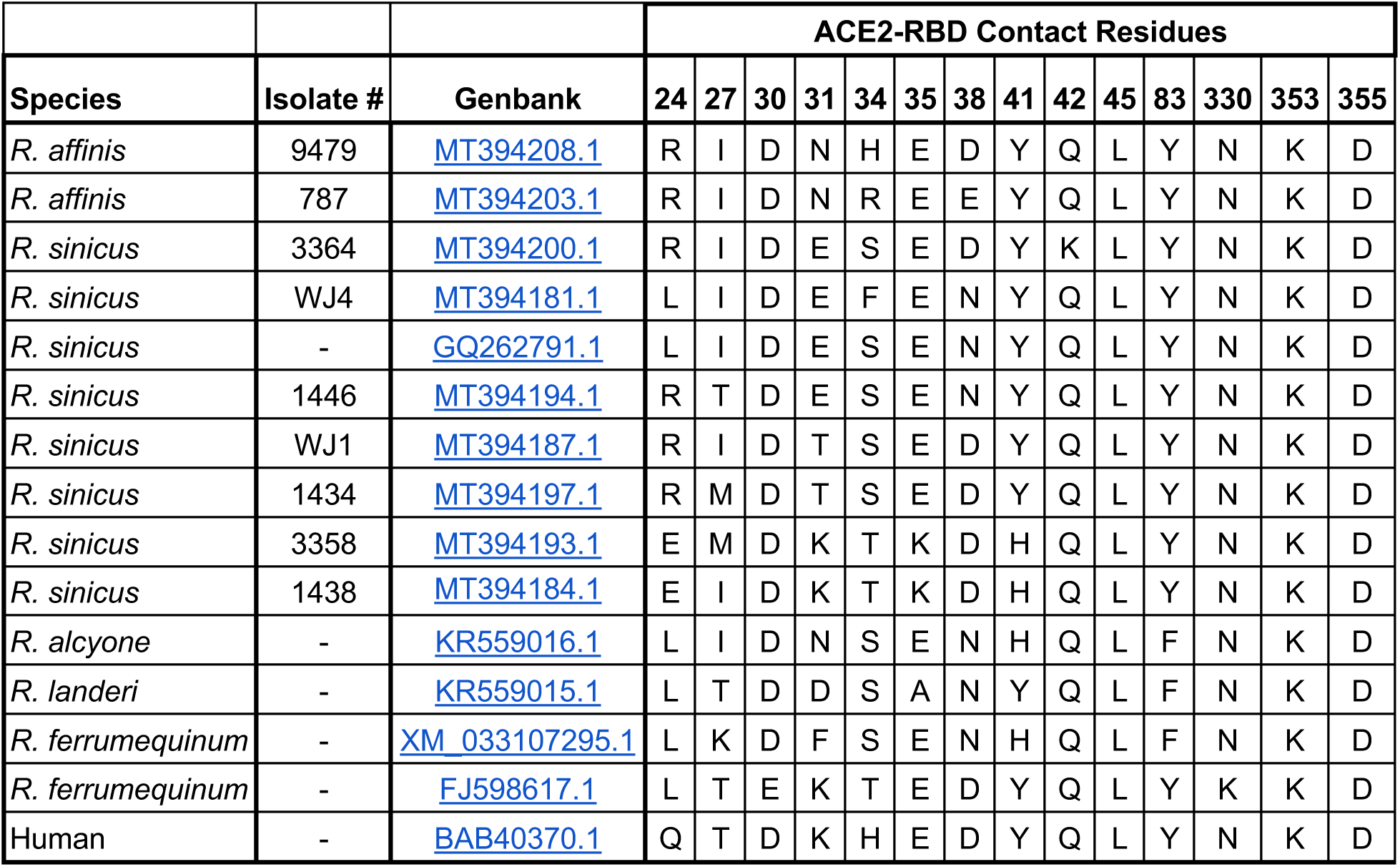
*Rhinolophus* ACE2 alleles and contact residues.

|  |  |  | ACE2-RBD Contact Residues |  |  |  |  |  |  |  |  |  |  |  |  |  |
| --- | --- | --- | --- | --- | --- | --- | --- | --- | --- | --- | --- | --- | --- | --- | --- | --- |
| Species | Isolate # | Genbank | 24 | 27 | 30 | 31 | 34 | 35 | 38 | 41 | 42 | 45 | 83 | 330 | 353 | 355 |
| *R. affinis* | 9479 | MT394208.1 | R | I | D | N | H | E | D | Y | Q | L | Y | N | K | D |
| *R. affinis* | 787 | MT394203.1 | R | I | D | N | R | E | E | Y | Q | L | Y | N | K | D |
| *R. sinicus* | 3364 | MT394200.1 | R | I | D | E | S | E | D | Y | K | L | Y | N | K | D |
| *R. sinicus* | WJ4 | MT394181.1 | L | I | D | E | F | E | N | Y | Q | L | Y | N | K | D |
| *R. sinicus* | - | GQ262791.1 | L | I | D | E | S | E | N | Y | Q | L | Y | N | K | D |
| *R. sinicus* | 1446 | MT394194.1 | R | T | D | E | S | E | N | Y | Q | L | Y | N | K | D |
| *R. sinicus* | WJ1 | MT394187.1 | R | I | D | T | S | E | D | Y | Q | L | Y | N | K | D |
| *R. sinicus* | 1434 | MT394197.1 | R | M | D | T | S | E | D | Y | Q | L | Y | N | K | D |
| *R. sinicus* | 3358 | MT394193.1 | E | M | D | K | T | K | D | H | Q | L | Y | N | K | D |
| *R. sinicus* | 1438 | MT394184.1 | E | I | D | K | T | K | D | H | Q | L | Y | N | K | D |
| *R. alcyone* | - | KR559016.1 | L | I | D | N | S | E | N | H | Q | L | F | N | K | D |
| *R. landeri* | - | KR559015.1 | L | T | D | D | S | A | N | Y | Q | L | F | N | K | D |
| *R. ferrumequinum* | - | XM_033107295.1 | L | K | D | F | S | E | N | H | Q | L | F | N | K | D |
| *R. ferrumequinum* | - | FJ598617.1 | L | T | E | K | T | E | D | Y | Q | L | Y | K | K | D |
| Human | - | BAB40370.1 | Q | T | D | K | H | E | D | Y | Q | L | Y | N | K | D |

**Table S2.** CryoEM data collection and refinement statistics.

|  | PRD-0038 RBD - *R. alcyone* ACE2<br>EMD-41786 | PRD-0038 RBD - *R. alcyone* ACE2<br>(Local refinement)<br>PDB 8U0T<br>EMD-41784 | PRD-0038 S<br>PDB 8U29<br>EMD-41842 | PRD-0038 S<br>(NTD local<br>refinement)<br>EMD-41843 |
| --- | --- | --- | --- | --- |
| Data collection and processing |  |  |  |  |
| Magnification | 105,000 | 105,000 | 105,000 | 105,000 |
| Voltage (kV) | 300 | 300 | 300 | 300 |
| Electron exposure (e<sup>-</sup>/Å<sup>2</sup>) | 60 | 60 | 60 | 60 |
| Defocus range (μm) | -0.2 - -3.0 | -0.2 - -3.0 | 0 - -3.0 | 0 - -3.0 |
| Pixel size (Å) | 0.843 | 0.843 | 0.843 | 0.843 |
| Symmetry imposed | C2 | C1 | C3 | C1 |
| Final particle images (no.) | 304,806 | 284,934 | 103,347 | 310,041 |
| Map resolution (Å) | 3.5 | 3.2 | 2.8 | 2.9 |
| FSC threshold | 0.143 | 0.143 | 0.143 | 0.143 |
| Map sharpening Bfactor (Å<sup>2</sup>) | -133 | -118 | -82.6 | -89.7 |
| Validation |  |  |  |  |
| MolProbity score |  | 1.08 | 1.28 |  |
| Clashscore |  | 1.62 | 1.69 |  |
| Poor rotamers (%) |  | 0.47 | 1.13 |  |
| Ramachandran plot |  |  |  |  |
| Favored (%) |  | 97.02 | 95.44 |  |
| Allowed (%) |  | 2.86 | 4.38 |  |
| Disallowed (%) |  | 0.12 | 0.19 |  |

**Table S3.** Representative K_D_ values determined from BLI binding analysis.

|  | K<sub>D</sub> (M) | K<sub>D</sub> Error | k<sub>a</sub> (1/Ms) | k<sub>a</sub> Error | k<sub>dis</sub> (1/s) | k<sub>dis</sub> Error |
| --- | --- | --- | --- | --- | --- | --- |
| PRD-0038 WT RBD<br>: monomeric *R. alcyone*<br>ACE2 | 5.07E-08 | 1.64E-10 | 1.64E+05 | 5.10E+02 | 8.33E-03 | 7.80E-06 |
| PRD-0038 T487W RBD<br>: monomeric *R. alcyone*<br>ACE2 | 2.12E-08 | 3.96E-11 | 1.24E+05 | 2.18E+02 | 2.62E-03 | 1.63E-06 |
| PRD-0038 WT RBD<br>: monomeric *R. landeri*<br>ACE2 | N.D. | N.D. | N.D. | N.D. | N.D. | N.D. |
| PRD-0038 T487W RBD<br>: monomeric *R. landeri*<br>ACE2 | 1.88E-07 | 4.22E-10 | 4.49E+04 | 9.64E+01 | 8.43E-03 | 5.62E-06 |

\* N.D. - not determined due to weak binding

## METHODS

### Cell lines

Cell lines used in this study were obtained from HEK293T (ATCC, CRL-11268) and Expi293F (Thermo Fisher Scientific, A145277) except for the HEK293T cells with stable human ACE2 expression which was kindly provided by Jesse Bloom^87^. Cells were cultured in 10% FBS, 1% penicillin-streptomycin DMEM at 37℃, 5% CO_2_. None of the cell lines were authenticated or tested for mycoplasma contamination.

### Production of recombinant PRD-0038 RBDs

The PRD-0038 S glycoprotein sequence was obtained from Genbank (<u>MT726045</u>). Both wildtype and mutant PRD-0038 RBDs (residues 318-520) were synthesized by GenScript with an N-terminal mu-phosphatase signal peptide and C-terminal 8x His tag, a short linker (GGSS) followed by an Avi tag in a pCMVR plasmid. Expi293F cells were grown at 37℃ with 8% CO_2_ and DNA transfections were conducted with the ExpiFectamine 293 Transfection Kit (Thermo Fisher Scientific). Cell culture supernatants were harvested three days post transfection. RBDs were purified using nickel based affinity chromatography using HisTrap™ High Performance column (Cytiva). Proteins were first washed with 10 column volumes of a buffer containing 25mM sodium phosphate (pH8.0) and 300 mM NaCl, before elution with 8 column volumes of a buffer containing 25 mM sodium phosphate (pH8.0), 300 mM NaCl, 500mM imidazole. Eluted proteins were buffer exchanged into 1x PBS (137mM NaCl, 2.7mM KCl, 10mM Na2HPO4, 1.8mM KH2PO4, pH 8.0) using Amicon Ultra-15 Centrifugal Filter Unit (10 kDa) (Millipore). Overnight biotinylation reactions were performed using the BirA Biotin-Protein Ligase Kit (Avidity) at 4℃ in 1x BiomixA, BiomixB. Biotinylated proteins were once again affinity purified using the HisTrap column as previously described to get rid of BirA. Once purified, buffer exchanged into PBS, and concentrated, proteins were flash-frozen and stored at –80℃ until use.

### Production of recombinant ACE2 ectodomains

Genbank accession numbers for all ACE2s can be found in **Table S1**. Recombinant ACE2 ectodomain constructs were synthesized by GenScript. ACE2-His ectodomain constructs comprise of residues 19–615 with an N-terminal mu-phosphatase signal peptide and C-terminal 8x His tag, a short GGSS linker, and an Avi tag. ACE2-Fc ectodomain constructs comprise residues 19–615 with an N-terminal mu-phosphatase signal peptide and C-terminal fusion to a sequence encoding a thrombin cleavage site, a short linker (GGGG) and a human Fc fragment and were cloned in a pCMV plasmid. The native *R. alcyone* ACE2 ectodomain dimer comprises residues 1-740 and a C-terminal 8x His tag, a GGSS linker and an Avi tag and was cloned in a pCMVR plasmid. All ACE2 orthologue ectodomains were produced in Expi293F cells at 37℃ supplemented and 8% CO_2_. Transfections were performed with the ExpiFectamine 293 Transfection Kit (Thermo Fisher Scientific). Cell culture supernatants were harvested four days after transfection and proteins were purified using HiTrap Protein A HP (Cytiva) or HisTrap™ High Performance column (Cytiva). ACE2-Fc proteins were first washed with 10 column volumes of 20 mM sodium phosphate (pH8.0) then eluted with 0.1 M citric acid (pH 3.0) directly into tubes containing 1M Tris-HCl (pH 9.0). Purified proteins were buffer exchanged into PBS (pH 8.0), concentrated using Spin-X® UF 20 mL Centrifugal Concentrator, 100,000 MWCO Membrane (PES) (Corning), and flash-frozen. ACE2-His proteins were washed with 10 column volumes of a buffer containing 25 mM sodium phosphate (pH 8.0), 300 mM NaCl, then eluted with 8 column volumes of a buffer containing 25 mM sodium phosphate (pH 8.0), 300 mM NaCl and 500 mM imidazole. Eluted proteins were buffer exchanged into 1x PBS (pH8.0) using Amicon Ultra-15 Centrifugal Filter Unit (10 kDa) (Millipore) and flash-frozen.

### Production of recombinant PRD-0038 PentaPro S, HexaPro S, and SARS-CoV-2 HexaPro S

The PRD-0038 S glycoprotein sequence was obtained from Genbank (<u>MT726045</u>). Recombinant PRD-0038 S glycoprotein ectodomain (residues 16-1194) constructs with pre-fusion stabilizing mutations (PentaPro: F800P, S882P, S925P, K969P, V970P or HexaPro: F800P, A875P, S882P, S925P, K969P, V970P) were synthesized by GenScript with an N-terminal mu-phosphatase signal peptide and C-terminal short linker (SG), TEV protease site (RENLYFQ), a short linker (GGGGSG), Foldon, 8x His tag, a short linker (GGSS) followed by an Avi tag in a pCMVR plasmid. The SARS-CoV-2 S glycoprotein ectodomain construct comprises residues 1-1208 with the native signal peptide, the HexaPro prefusion stabilizing mutations^56^ (F817P, A892P, A899P, A942P, K986P, V987P), abrogation of the furin cleavage site (residues 682-685, GSAS) and a C-terminal short linker (GSG), followed by a foldon, HRV 3C site (LEVLFQGP), a short linker (GSG), an avi tag, a short linker (GSG), an 8x his tag in a pcDNA3.1(-) plasmid. Expi293F cells were grown at 37℃ with 8% CO_2_ and DNA transfections were conducted with the ExpiFectamine 293 Transfection Kit (Thermo Fisher Scientific). Cell culture supernatants were harvested four days post-transfection and proteins were purified using HisTrap™ High Performance column (Cytiva). Proteins were first washed with 10-15 column volumes of a buffer containing 25 mM sodium phosphate, 300 mM NaCl, 20 mM imidazole, pH 8.0, followed by elution with 10-15 column volumes using 300 mM imidazole, pH 8.0. Eluted proteins were concentrated and buffer exchanged into 1x PBS or 1x TBS (20 mM Tris, 150 mM NaCl, pH 8.0) using Amicon Ultra-15 Centrifugal Filter Unit (100 kDa) (Millipore). Purified proteins were snap frozen and stored at – 80°C. Purified proteins were checked for endotoxin level using Charles River Limulus Amebocyte Lysate (LAL) cartridges PTS201F. Endotoxin-free SARS-CoV-2 HexaPro S and PRD-0038 HexaPro S were flash-frozen and stored at –80°C until the day of immunization.

### Production of PRD-0038 wild-type RBD-Natively Dimerized R. alcyone Complex

For complex formation, wild-type PRD-0038 RBD was mixed with natively dimerized R. alcyone ACE2 His at a 4:1 molar ratio, then incubated at room temperature for 5-10 min. Gel filtration was performed to remove excess RBD on a Superose 6 10/300 GL column (Cytiva) equilibrated in 50 mM Tris-HCl, 150 mM NaCl. Complex formation was confirmed by SDS-PAGE, and the PRD-0038 RBD-R. alcyone ACE2 complex was snap frozen and stored at –80°C until day of grid preparation.

### Binding analysis using biolayer interferometry (BLI)

BLI binding assays were performed on an Octet Red (Sartorius) instrument operated at 30℃ with shaking (1000 rpm). For biotinylated RBD and ACE2-Fc or ACE2-His binding assays, streptavidin biosensors were hydrated in water for 10 min prior to the experiment. Biosensors were incubated in 10x kinetics buffer for 60s followed by the loading of biotinylated RBDs to the tip, all to a final level of 1 nm. Loaded biosensors were equilibrated in 10x kinetics buffer (Sartorius) for 120s which served as our baseline. For avidity binding assays, association with 1 µM ACE2-Fc (dimeric form) was performed for 300 s followed by 300 s of dissociation in 10x kinetics buffer. For affinity binding assays to determine K_D_ values, RBD-loaded tips were dipped into a concentration series of ACE2-His (2 fold serial dilution from 100 nM to 6.25 nM for *R. alcyone* ACE2-His, 3 fold serial dilution from 660 nM to 8 nM for *R. landeri* ACE2-His) for 600 s followed by 600 s of dissociation in 10x kinetics buffer. Global fits were used to calculate K_D_ values using a 1:1 binding fit model. Data were plotted using GraphPad Prism. Assays were replicated with three biological replicates (recombinant RBD proteins generated on different days) and representative graphs are shown.

### Production of VSV pseudoviruses

Wildtype and mutant PRD-0038 S constructs consisting of residues 1-1235 and containing a 21 residue C-terminal deletion (del21) followed by a 3x FLAG tag were synthesized by GenScript and placed into an HDM plasmid. VSV pseudoviruses were produced using HEK293T cells seeded on BioCoat Cell Culture Dish: poly-D-Lysine 100 mm (Corning). Cells were transfected with PRD-0038 S-Flag constructs using Lipofectamine 2000 (Life Technologies) in Opti-MEM transfection medium. After 5h of incubation at 37 °C with 5% CO2, cells were supplemented with DMEM containing 10% of FBS. On the next day, cells were infected with VSV (G*ΔG-luciferase) for 2h, followed by five time wash with DMEM medium before addition of anti-VSV G antibody (I1-mouse hybridoma supernatant diluted 1:40, ATCC CRL-2700) and medium. After 18-24 h of incubation at 37 °C with 5% CO_2_, pseudoviruses were collected and cell debris removed by centrifugation at 3,000xg for 10 min. Pseudoviruses were further filtered using a 0.45 µm syringe filter and concentrated 25-50x prior to storage at –80°C. Mock VSV pseudoviruses were prepared as above but without S transfection.

### Cell entry assays comparing wildtype and mutant PRD-0038 S VSV pseudoviruses

HEK293T cells were transfected with full length *Rhinolophus* ACE2 orthologs using Lipofectamine 2000 (Life Technologies) in Opti-MEM five hours prior to plating into 96-well plates [3610] (Corning) coated with poly-lysine [P4707] (Sigma) and incubated 18-24 h before infection with VSV pseudoviruses. For human ACE2 entry assays, HEK293T cells with stable hACE2 expression were plated into poly-lysine-coated 96-well plates and incubated for 18-24h before infection with VSV pseudoviruses. The amount of pseudovirus used for infection was adjusted using Western Blot based on S incorporation across different mutants to use a constant input of S. Detection of VSV backbone was performed with 1:1,000 Anti-VSV-M [23H12] Antibody (Kerafast). Detection of 3x-FLAG tagged S was performed with 1:400 monoclonal ANTI-FLAG® M2 antibody [F3165] produced in mouse (Sigma). 1:50,000 Alexa Fluor® 680 AffiniPure Goat Anti-Mouse IgG [115-625-174] (Jackson ImmunoResearch) was used as the secondary antibody. A representative Western Blot is shown in Figure S4E. Genbank accession numbers for all ACE2s can be found in Table S1. After 1h of infection, an additional 40 μL of DMEM supplemented with 20% FBS and 2% PenStrep was added to the cells. After 18–20h, 40 μL of One-Glo-EX substrate (Promega) was added to each well and incubated on a plate shaker in the dark for 5 min before reading the relative luciferase units using a BioTek Neo2 plate reader. Fold change of relative luciferase units over mock VSV were plotted in Prism (GraphPad) with mock being cells that were not transfected with an S-encoding plasmid. 3 biological replicates, each of which with 3 technical replicates were carried out.

### Cell entry assay for PRD-0038 S VSV with distinct *Rhinolophus* ACE2s

For wildtype PRD-0038 S VSV entry into HEK293T cells transfected with *R. affinis*, *R. sinicus, R. landeri,* and *R. alcyone* alleles full-length ACE2 alleles (Figure 1D), HEK293T cells (ATCC) were cultured in 10% FBS, 1% penicillin–streptomycin DMEM at 37 °C in a humidified 5% CO_2_ incubator. Cells were plated 18-24 hours prior to transfection into 96-well plates [3610] (Corning) coated with poly-lysine [P4707] (Sigma). All transfections were performed using full-length *Rhinolophus* ACE2 placed into a HDM plasmid (synthesized by GenScript). Transfection of ACE2 alleles into HEK293T cells was performed using 0.2 µg DNA and 0.15 µL Lipofectamine 2000 (Life Technologies) per well in Opti-MEM. After a 5 h incubation at 37 °C in a humidified 8% CO_2_ incubator, DMEM was added to obtain a final concentration of 10% FBS and 1% penicillin– streptomycin. Cells were incubated at 37 °C in a humidified 8% CO_2_ incubator for 36–48 h prior to infection. For each infection test, 2-3 technical replicates were performed, and the assays were repeated on a second day, for a total of 4-6 technical replicates. Three biological replicates (pseudovirus generated on different days) were used for cell entry and each point shown represents the mean fold change for each biological replicate. Results were plotted using Graphpad Prism (Figure 1D). Genbank accession numbers for all ACE2s can be found in Table S1.

### Cryo-EM sample preparation and data collection

Cryo-EM grids of PRD-0038 PentaPro S were prepared using two separate methods and data were combined during data processing. The first dataset was collected from the grids prepared using a reverse grid-blotting method. 3 µL of sample was added to the carbon side of a glow discharged C-flat R2/2 copper grid and 1µL was added to the back side before addition of 1 µL of CHEMS lipid dissolved in chloroform on the front side. The sample was allowed to sit on the grid for 1 minute and then manually blotted from the back side using a strip of Whatman #1 filter paper and plunged into liquid ethane. The second dataset was collected from a lacey carbon grid with a thin home-made continuous carbon layer. 3 µL of 0.15 mg/mL PRD-0038 Pentapro S was loaded onto the glow discharged (6s at 20mA) grid followed by plunge freezing using a vitrobot MarkIV (ThermoFisher Scientific). The grid was blotted with a blot force of –1, 3 second blot time, and 10 second wait time before the plunge freeze at 100% humidity and 25 °C. For the PRD-0038 RBD-ACE2 complex, grids were prepared by applying 3 µL of 4 mg/ml PRD-0038 RBD bound to the *R. alcyone* ACE2 dimer with 7 mM CHAPSO (Anatrace) were applied and blotted twice as previously described^88^, onto freshly glow discharged R 2/2 UltrAuFoil grids prior to plunge freezing using a vitrobot MarkIV (ThermoFisher Scientific) with a blot force of 0 and 5 sec blot time at 100 % humidity and 22°C. Data were acquired using an FEI Titan Krios transmission electron microscope operated at 300 kV and equipped with a Gatan K3 direct detector and Gatan Quantum GIF energy filter, operated in zero-loss mode with a slit width of 20 eV. Automated data collection was carried out using Leginon^89^ at a nominal magnification of 105,000x with a pixel size of 0.843 Å. The dose rate was adjusted to 15 counts/pixel/s, and each movie was acquired in counting mode fractionated in 75 frames of 40 ms. A total of 2,482 and 11,743 micrographs were collected for the PRD-0038 S and PRD-0038 RBD-ACE2 datasets, respectively.

### Cryo-EM data processing, model building and refinement

Motion correction, contrast-transfer function (CTF) parameter estimation, automatic particle picking, and extraction were performed using Warp^90^ for each data set. For the PRD-0038 S structure, particle images were extracted with a box size of 260 pixels with a pixel size of 1.686Å. After two rounds of 2D classification using cryoSPARC^91^, well-defined particles were selected and particles from each dataset were combined and binned to a box size of 130 pixels with a pixel size of 3.372Å for subsequent 3D classification using Relion^92,93^ with 50 iterations (angular sampling 7.5° for 25 iterations and 1.8° with local search for 25 iterations). 103,347 particles were selected and re-extracted with a box size of 260 and pixel size of 1.686Å for cryoSPARC non-uniform refinement^94^ with C3 symmetry and further subjected to Bayesian polishing^95^ in Relion. Finally, another round of non-uniform refinement with C3 symmetry and optimized per-particle defocus was carried out to the polished particles. To improve the density of the NTD, we used symmetry expansion and local refinement using cryoSPARC. For the PRD-0038 RBD – *R. alcyone* ACE2 structure, two rounds of reference-free 2D classification were performed using cryoSPARC to select well-defined particle images. These selected particles were subjected to two rounds of 3D classification with 50 iterations each (angular sampling 7.5° for 25 iterations and 1.8° with local search for 25 iterations) using Relion with an initial model generated with ab-initio reconstruction in cryoSPARC. 3D refinements were carried out using non-uniform refinement along with per-particle defocus refinement in CryoSPARC. To improve the density of the RBD-ACE2 dimer, the particles were subjected to cryoSPARC heterogeneous refinement. Particles belonging to classes with the best resolved RBD-ACE2 density were selected and subjected to the Bayesian polishing procedure implemented in Relion before performing another round of non-uniform refinement in cryoSPARC followed by per-particle defocus refinement and again non-uniform refinement. To further improve the density of the RBD-ACE2 domains, the particles were symmetry expanded and subjected to focus 3D classification without refining angles and shifts using a soft mask encompassing the RBD and monomer ACE2 using a tau value of 40 in Relion. Particles belonging to classes with the best resolved RBD-ACE2 density were selected and then subjected to local refinement using CryoSPARC. Local resolution estimation, filtering, and sharpening were carried out using CryoSPARC. Reported resolutions are based on the gold-standard Fourier shell correlation (FSC) of 0.143 criterion and Fourier shell correlation curves were corrected for the effects of soft masking by high-resolution noise substitution^96,97^. UCSF Chimera and Coot were used to fit atomic models into the cryoEM maps. S and RBD-ACE2 models were refined and relaxed using Rosetta^98,99^ using sharpened and unsharpened maps and validated using Phenix^100^, Molprobity^101^ and Privateer^102^.

### Monoclonal antibody ELISAs

For PRD-0038 HexaPro S and SARS-CoV-2 HexaPro S ELISAs, 30 μl of the proteins at 3 μg/mL were plated onto 384-well Nunc Maxisorp plate (ThermoFisher, 464718) in 1x TBS and incubated 1h at 37°C followed by slap drying and blocking with 80 μL of Casein for 1 h at 37°C. After incubation, plates were slap dried and 1:4 serial dilutions of the corresponding mAbs starting from 0.1 mg/ml were made in 30 μl TBST, added to the plate and incubated at 37°C for 1 h. Plates were washed 4x in TBST and 30 μl of 1:5,000 Goat anti-Human IgG Fc Secondary Antibody, HRP (Thermo Fisher, A18817) or Goat anti-Syrian Hamster IgG (H+L) Secondary Antibody, HRP (Thermo Fisher, PA1-28823) were added to each well and incubated at 37°C. After 1 h, plates were washed 4x in TBST and 30 μl of TMB (SeraCare) was added to every well for 2 min at room temperature. Reactions were quenched with the addition of 30 μl of 1N HCl. Plates were immediately read at 450 nm on a BioTek Neo2 plate reader and data plotted and fit in Prism 9 (GraphPad) using nonlinear regression sigmoidal, 4PL, X is the concentration to determine EC_50_ values from curve fits.

### Deep mutational scanning for mutational effects on ACE2 binding

The complete deep mutational scanning pipeline can be found at: https://github.com/tstarrlab/SARSr-CoV-RBD_DMS/blob/main/results/summary/summary.md. Deep mutational scanning libraries for sarbecovirus strains including PRD-0038 and SARS-Cov-1 Urbani were constructed as previously described^76^. Briefly, site-saturation mutagenesis libraries spanning all RBD positions were produced by Twist Bioscience (or NNS mutagenesis for positions that failed Twist mutagenesis), tagged with an N16 barcode, and cloned into a yeast display vector backbone via Gibson Assembly. Libraries were electroporated into *E. coli* and plated at a target bottleneck of 40,000 unique barcodes per library to overrepresent the ∼4,000 possible amino acid mutations. Colonies were scraped from each transformation plate, library plasmid purified, and transformed into the AWY101 *S. cerevisiae* strain^103^ for yeast surface-display experiments. Library plasmids were sequenced using a PacBio Sequel IIe to generate long sequence reads spanning the N16 barcodes and RBD coding sequence. Raw CCS reads are available on the NCBI Sequence Read Archive, BioProject PRJNA962117, BioSample SAMN34384156. Reads were processed using alignparse (version 0.2.4)^104^ to generate a table linking each N16 barcode to its unique RBD mutant, available at: https://github.com/tstarrlab/SARSr-CoV-RBD_DMS/blob/main/results/variants/codon_variant_table_PRD0038.csv and https://github.com/tstarrlab/SARSr-CoV-RBD_DMS/blob/main/results/variants/codon_variant_table_SARS1.csv.

The RBD expression level and ACE2-binding avidity of each RBD mutant was determined via high-throughput FACS-seq assays as previously described^76^. ACE2-binding titrations were performed by incubating induced yeast-display libraries with a concentration series of dimeric ACE2 ligands from 10^-6^ to 10^-13^ M at 1-log intervals, plus a 0 M ACE2 sample, with samples equilibrated overnight at room temperature with mixing. Yeast were washed with PBS-BSA (0.2 mg/L), labeled with 1:100 FITC-conjugated chicken anti-Myc (Immunology Consultants CMYC-45F) to detect yeast-displayed RBD and 1:200 PE-conjugated streptavidin (Thermo Fisher S866) or goat anti-human-IgG (Jackson ImmunoResearch 109-115-098) to detect binding of biotinylated (human, Acro Biosystems H82E6) or Fc-tagged (*R. alcyone, R. landeri*) ACE2. For each titration sample, RBD^+^ yeast were fractionated into four bins of PE fluorescence (ACE2 binding), grown overnight, plasmid isolated, N16 barcode amplified, and barcodes counted via high-throughput sequencing on an Illumina NextSeq. RBD expression was measured by sorting cells into four bins on the basis of Myc-FITC labeling, followed by outgrowth, plasmid isolation, N16 barcode amplification, and sequencing. Sequencing reads are available on the NCBI Sequence Read Archive, BioProject PRJNA962117, BioSample SAMN34384823.

Demultiplexed Illumina barcode reads were aligned to library barcodes using dms_variants (version 0.8.9), yielding a table of counts of each barcode in each FACS bin which is available at https://github.com/tstarrlab/SARSr-CoV-RBD_DMS/blob/main/results/counts/variant_counts.csv.gz. Reads were downweighted by the ratio of total sequence reads from a bin to the number of cells sorted into that bin. For each barcode, we inferred the apparent dissociation constant for avid binding (K_D,app_) by fitting the standard non-cooperative Hill equation to the mean FACS bin of a barcode variant as a function of ACE2 concentration. For each barcode, expression was determined via a maximum likelihood estimator of log-MFI based on the distribution of barcode counts across FACS bins and the known fluorescence boundaries of those bins. The computational pipelines for computing per-barcode binding constants and expression phenotypes are available at: https://github.com/tstarrlab/SARSr-CoV-RBD_DMS/blob/main/results/summary/compute_binding_Kd_huACE2.md and https://github.com/tstarrlab/SARSr-CoV-RBD_DMS/blob/main/results/summary/compute_expression_meanF.md. Because most mutants in the library were independently associated with more than one N16 barcode, we derived the final mutant phenotype as the average of per-barcode measurements, as computed at: https://github.com/tstarrlab/SARSr-CoV-RBD_DMS/blob/main/results/summary/collapse_barcodes_lib40_41.md. The final per-mutant deep mutational scanning phenotypes are available at: https://github.com/tstarrlab/SARSr-CoV-RBD_DMS/blob/main/results/final_variant_scores/final_variant_scores_lib40_41.csv.

### Immunogenicity

Female BALB/c mice were purchased from Envigo (order code 047) at 7 weeks of age and were maintained in a specific pathogen-free facility within the Department of Comparative Medicine at the University of Washington, Seattle, accredited by the Association for Assessment and Accreditation of Laboratory Animal Care (AAALAC). Prior to each immunization, immunogens (endotoxin-free SARS-CoV-2 HexaPro S or PRD-0038 HexaPro S) were diluted to 20 µg/mL in 1x TBS (20mM Tris, 150mM NaCl, pH 8.0) and mixed with 1:1 vol/vol AddaVax (InvivoGen vac-adx-10) to reach a final dose of 1 µg of immunogen per injection. At 8 weeks of age, 6 mice per group were anesthetized and injected intramuscularly in the quadriceps with 50µL of immunogen per leg, 100µL total at weeks 0, 3, and 6. Mice were bled via the submental route at weeks 0, 2, 5, and 8. Blood was collected in serum separator tubes (BD # 365967) and rested for 30 min at room temperature for coagulation. Serum tubes were then centrifuged for 10 min at 2,000 x g and serum was collected and stored at –80°C until use. Animal experiments were conducted in accordance with the University of Washington’s Institutional Animal Care and Use Committee.

### Neutralization assays

For mAb neutralization against SARS-CoV-2 S VSV and PRD-0038 S K482Y/T487W VSV, HEK293T cells with stable human ACE2 expression in DMEM supplemented with 10% FBS and 1% PenStrep were seeded at 40,000 cells/well into 96-well plates [3610] (Corning) coated with poly-lysine [P4707] (Sigma) and incubated overnight at 37°C. After 16-20h of incubation, a half-area 96-well plate (Greiner) was prepared with 1:5 serial dilutions of S2X259 and S2X35 starting from 0.1 mg/ml in DMEM, and 1:3 serial dilutions of 76E1 starting from 0.15 mg/ml in DMEM, for a total of 22 µL per well. An equal volume of DMEM with diluted pseudoviruses was added to each well. All pseudoviruses were diluted between 1:3-1:27 to reach a target entry of ∼10^6^ RLU. The mixture was incubated at room temperature for 45-60 minutes. Media was removed from the cells and 40 μL from each well of the half-area 96-well plate containing mAb and pseudovirus were transferred to the 96-well plate seeded with cells and incubated at 37°C for 1h. After 1h, an additional 40 μL of DMEM supplemented with 20% FBS and 2% PenStrep was added to the cells. After 18–20h, 40 μL of One-Glo-EX substrate (Promega) was added to each well and incubated on a plate shaker in the dark for 5 min before reading the relative luciferase units using a BioTek Neo2 plate reader. Relative luciferase units were plotted and normalized in Prism (GraphPad): 100% neutralization being cells lacking pseudovirus and 0% neutralizing being cells containing virus but lacking mAb. Prism (GraphPad) nonlinear regression with “[inhibitor] versus normalized response with a variable slope” was used to determine IC_50_ values from curve fits with 3 technical repeats. 3 biological replicates were carried out for each mAb.

For SARS-CoV-2 D614G S VSV, BA.2 S VSV, BA.5 S VSV, RaTG13 S VSV, and SARS-CoV-1 S VSV neutralization, HEK293T cells with stable human ACE2 expression in DMEM supplemented with 10% FBS and 1% PenStrep were seeded at 40,000 cells/well into 96-well plates [3610] (Corning) coated with poly-lysine [P4707] (Sigma) and incubated overnight at 37°C. For PRD-0038 S VSV and Khosta1 S VSV neutralization, HEK293T cells were transfected with full length *R. alcyone* ACE2 using Lipofectamine 2000 (Life Technologies) in Opti-MEM five hours prior to plating into 96-well plates [3610] (Corning) coated with poly-lysine [P4707] (Sigma) and incubated overnight at 37°C. The following day, a half-area 96-well plate (Greiner) was prepared with 3-fold serial sera dilutions (starting dilutions determined for each serum and pseudovirus, 22uL per well). An equal volume of DMEM with diluted pseudoviruses was added to each well. All pseudoviruses were diluted between 1:3-1:27 to reach a target entry of ∼10^6^ RLU. The mixture was incubated at room temperature for 45-60 minutes. Media was removed from the cells and 40 μL from each well of the half-area 96-well plate containing sera and pseudovirus were transferred to the 96-well plate seeded with cells and incubated at 37°C for 1h. After 1h, an additional 40 μL of DMEM supplemented with 20% FBS and 2% PenStrep was added to the cells. After 18–20h, 40 μL of One-Glo-EX substrate (Promega) was added to each well and incubated on a plate shaker in the dark for 5 min before reading the relative luciferase units using a BioTek Neo2 plate reader. Relative luciferase units were plotted and normalized in Prism (GraphPad): 100% neutralization being cells lacking pseudovirus and 0% neutralizing being cells containing virus but lacking sera. Prism (GraphPad) nonlinear regression with “log[inhibitor] versus normalized response with a variable slope” was used to determine ID_50_ values from curve fits with 3 technical repeats. 3 biological replicates were carried out for each sample-pseudovirus pair.

### Breadth– and epitope-mapping of vaccine sera via deep mutational scanning

The complete serum deep mutational scanning pipeline is described at: https://github.com/tstarrlab/SARSr-CoV_MAP_PRD0038-vaccine/blob/main/results/summary/summary.md. Binding of serum was evaluated against deep mutational scanning pools for PRD-0038 and SARS-CoV-1 whose construction is described above, previously published deep mutational scanning pools for SARS-CoV-2 Wuhan-Hu-1^76^ and Omicron BA.2^38^, and a previously published pan-sarbecovirus panel^29^ that was supplemented with additional newly described sarbecovirus and SARS-CoV-2 variants. Serum was first depleted of non-specific yeast-reactive antibodies as previously described^105^. Yeast-display RBD libraries were pooled, induced for yeast surface expression, and labeled with serum at 1:100, 1:1000, 1:10,000, and 1:100,000 dilutions for one hour at room temperature. Yeast were washed with PBS-BSA and labeled with secondary Myc-FITC antibody and APC-conjugated goat anti-mouse-IgG (Jackson ImmunoResearch 115-605-008). As with ACE2-binding titrations, libraries were then partitioned into four bins of serum binding on a BD FACSAria, collecting a minimum of 6 million RBD^+^ cells per sample concentration across the four bins. Cells were grown post-sort, plasmid purified, N16 barcode amplified, and sequenced on an Illumina NextSeq. Raw Illumina sequencing data is available from the NCBI Sequence Read Archive, BioProject PRJNA714677, BioSample SAMN36715819. Barcode reads were mapped to library barcodes, with raw counts found at: https://github.com/tstarrlab/SARSr-CoV_MAP_PRD0038-vaccine/blob/main/results/counts/variant_counts.csv.

For each library barcode, an area under the curve (AUC) metric was derived from its distribution of sequence reads across sort bins. First, the strength of serum binding to each barcode at each serum dilution was determined as the simple mean bin from cell counts across integer-weighted bins, and subtracted by background mean bin determined from a sort from yeast libraries not incubated with sera. Any barcode with less than 3 cell counts at any sample concentration was eliminated from analysis. An AUC metric was then calculated from the relationship between mean bin and serum dilution. AUC calculation can be found at: https://github.com/tstarrlab/SARSr-CoV_MAP_PRD0038-vaccine/blob/main/results/summary/compute_AUC.md, and per-barcode AUC metrics are available at: https://github.com/tstarrlab/SARSr-CoV_MAP_PRD0038-vaccine/blob/main/results/bc_sera_binding/bc_sera_binding.csv. We then computed the per-variant AUC as the robust mean of replicate barcodes linked with the identical RBD variant, by taking the mean per-barcode AUC after trimming tails of the top and bottom 2.5% of AUC values among the replicate barcodes. Because mutations that disrupt RBD expression artifactually decrease serum binding, we applied two final filters: first, we censored the AUC measurement for any mutant with a measured impact on RBD expression of greater than one log-MFI unit (RBD expression < –1) from DMS measurements described above; and second, we derived a normalization constant from the slope of the linear model relating serum AUC and expression globally across all library variants for variants with expression >-1, and normalized our raw AUC measurements by this constant. The final variant derivation can be found at: https://github.com/tstarrlab/SARSr-CoV_MAP_PRD0038-vaccine/blob/main/results/summary/collapse_barcodes_SARSr-DMS.md, and final per-variant serum-binding values are available at: https://github.com/tstarrlab/SARSr-CoV_MAP_PRD0038-vaccine/blob/main/results/final_variant_scores/final_variant_scores_wts.csv and https://github.com/tstarrlab/SARSr-CoV_MAP_PRD0038-vaccine/blob/main/results/final_variant_scores/final_variant_scores_dms.csv.

